# Müllerian duct maintenance at the cranial region is promoted by *Gata2*

**DOI:** 10.64898/2026.03.11.711111

**Authors:** Shuai Jia, Allyssa Fogarty, Wenyan Bai, Fei Zhao

## Abstract

The paradigm of sexual differentiation holds that female embryos retain Müllerian ducts (the precursor to female reproductive tract) by default in the absence of anti-Müllerian hormone (AMH). However, whether Müllerian duct maintenance requires active signaling has remained unclear. Here, we discover that the deletion of mesenchymal *Gata2* induces selective regression of the cranial Müllerian duct (future oviduct). This regression is not driven by ectopic AMH signaling, but rather by the loss of region-specific NRG1 signaling. In contrast, the caudal Müllerian duct (future uterus) is retained and displays disrupted epithelial differentiation upon *Gata2* deletion in either the mesenchymal or epithelial compartments. Our findings reveal a region-specific, GATA2-dependent program that actively maintains the cranial Müllerian duct, reshaping our understanding of female reproductive tract development.

## Main Text

A mammalian embryo initially forms both the primitive male and female reproductive tracts, which are known as the Wolffian duct (WD) and Müllerian duct (MD), respectively. During sexual differentiation, the embryo eliminates one of these primitive ducts, retaining only the WD in males and the MD in females (*1*). Jost’s landmark experiments in mid-20^th^ century revealed that this dimorphic process is driven by two testis-derived hormones: the testosterone that promotes WD differentiation, and a second substance then termed as the “Müllerian inhibitor” responsible for MD regression (*2, 3*). Subsequent molecular studies identified this Müllerian-inhibiting substance as anti-Müllerian hormone (AMH), a member of the transforming growth factor β family (*4–6*). Loss-of-function mutations in *Amh* lead to persistent Müllerian duct maintenance in males, whereas ectopic AMH signaling induces MD regression in females (*7*). These findings established the classical paradigm of sexual differentiation: female embryos maintain MDs due to the absence of AMH, which is therefore considered as a passive process.

The fate and differentiation of MDs rely on paracrine signals from the surrounding undifferentiated connective tissue known as the mesenchyme. The AMH-specific receptor, anti-Müllerian hormone receptor type II (AMHR2), is expressed in MD mesenchyme, where it activates R-SMADs and SP7 transcriptional effectors to induce MD regression in males (*8, 9*). In females, region-specific mesenchymal cues dictate MD differentiation into functionally and morphologically distinct organs along the craniocaudal axis, the oviduct, uterus, cervix and upper vagina (*10*). For instance, vaginal epithelium combined with the mesenchyme from the neonatal uterus adopted uterine morphology, whereas uterine epithelium grown underlying the vaginal mesenchyme underwent vaginal differentiation (*11*). The MD mesenchyme expresses HOXA9-13 genes, which provide the positional identity to epithelial cells along the craniocaudal axis (*12*). Collectively, these studies underscore the pivotal role of the mesenchyme in the fate decision and differentiation of MD epithelium.

GATA2, a member of the GATA transcription factor family, plays crucial roles in the development of multiple organs (*13–15*). It is expressed in the mesenchyme of urogenital organs, including the MD (*16–18*). Germline deletion of *Gata2* results in embryonic lethality at embryonic day (E) 10.5 due to defective hematopoiesis in mice (*19*), precluding investigation of its specific role in fetal MD development. Here, we utilized tissue-specific conditional knockout models to uncover the region-specific functions of mesenchymal *Gata2* in MD maintenance and differentiation in female embryos.

### Regression of the cranial MD in *Gata2^cKO^* female embryos

GATA2 expression was detected in the nuclei of mesenchymal cells and coelomic epithelium in both the cranial and caudal regions at E12.5 (the time of MD elongation), between E14.5-E16.5 (MD regression in males and maintenance in females), and postnatal day 0 (PND0, birth) (Fig. 1A and fig. S1, A and B). In the cranial region that gives rise to the oviduct, GATA2 expression was limited to the mesenchyme. However, in the caudal region that will differentiate into the uterus, GATA2 expression was observed in some MD epithelial cells at E16.5 and PND0. The dynamic expression of GATA2 in the cranial and caudal MD suggests that GATA2 could play cell type-specific and region-specific roles in MD differentiation.

**Fig. 1.**
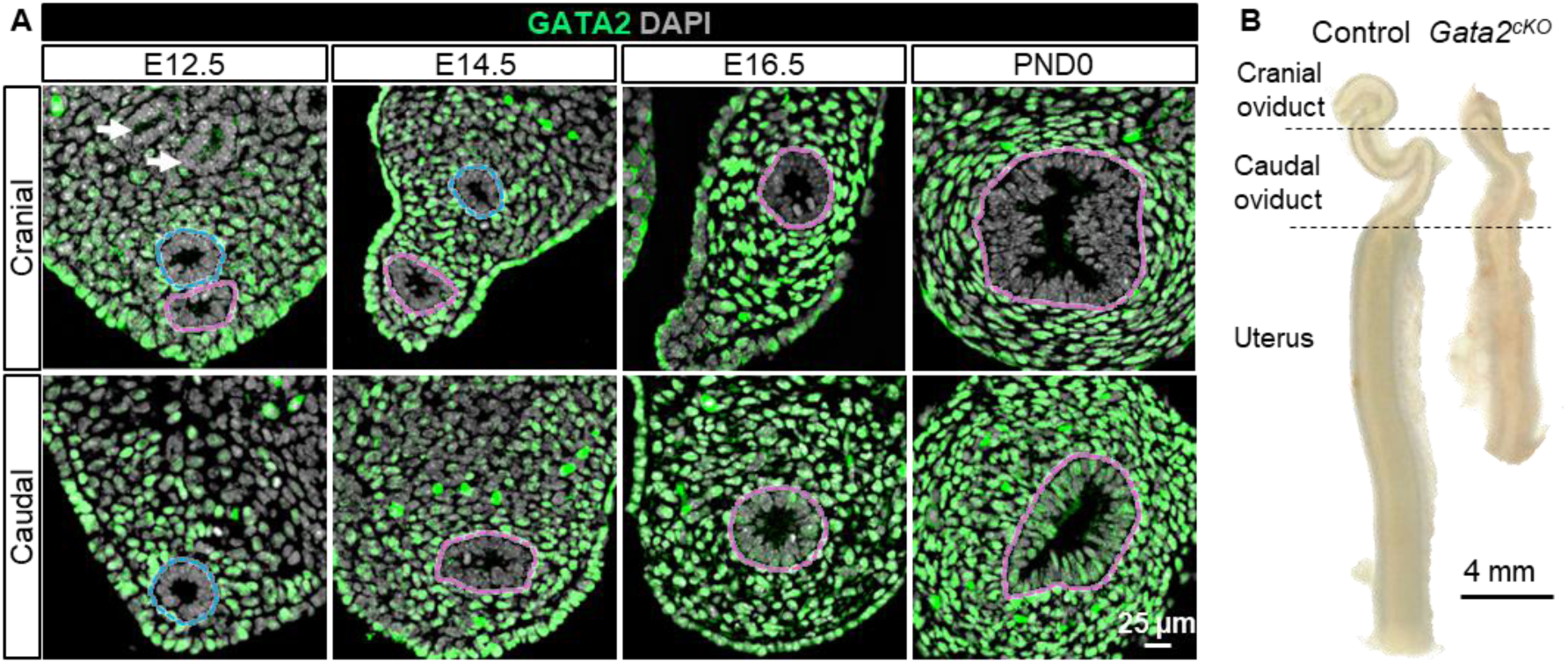
Gata2cKO driven by *Osr2-Cre* caused Müllerian duct hypoplasia and the loss of the cranial oviduct in female embryos. (**A**) GATA2 immunofluorescent staining in cranial and caudal mesonephroi at E12.5 (*n*=3), E14.5 (*n*=4), E16.5 (*n*=8) and PND0 (*n*=5). Blue and pink dashed lines: Wolffian ducts and Müllerian ducts, respectively; White arrows: mesonephric tubules. (**B**) Bright-field images of control and *Gata2^cKO^* upper female reproductive tract at PND0. *n*=3 per group.

To investigate the function of mesenchymal *Gata2*, we generated conditional *Gata2* knockout model (*Osr2^Cre+^:Gata2^f/f^*, *Gata2^cKO^*) with knock-in *Osr2-Cre* line that targets the mesonephric mesenchyme (*20, 21*). *Osr2* was expressed in the mesonephric mesenchyme around E10.5 (*22*). By crossing with the *Rosa-tdTomato* fluorescent reporter, we confirmed *Osr2-Cre* activity in the MD mesenchyme at E12.5, E14.5 and PND0 (fig. S2A). The tdTomato+ cells were also observed in some MD epithelial cells, suggesting that these cells might result from mesenchymal-to-epithelial transition during MD development (*23*). In *Gata2^cKO^* mesonephroi, *Gata2* expression was significantly reduced (fig. S2B). Collectively, these results demonstrate the establishment of the conditional deletion of *Gata2* in *Osr2*+ mesenchymal lineage.

During fetal development, the upper female reproductive tract elongates and differentiates into three morphologically distinct sections: the cranial oviducts, the caudal oviducts with a narrower lumen, and the uteri (Fig. 1B). By contrast, the upper female reproductive tract in *Gata2^cKO^* embryos was shortened and hypoplastic (Fig. 1B and fig. S3, A and B) with decreased cell proliferation (fig. S4, A and B) and increased cell apoptosis (fig. S4, C and D). More strikingly, *Gata2^cKO^* embryos lacked the cranial oviduct, the precursor to the infundibulum and adjacent ampulla regions that are essential for oocyte pickup and transport. At adulthood, the retained oviducts were blunted and fluid-filled due to the loss of cranial portion (fig. S3D). Taken together, our results demonstrate that mesenchymal *Gata2* plays a crucial role in promoting MD growth and maintaining the cranial MD.

### No ectopic AMH/AMHR2 signaling in *Gata2^cKO^* female mesonephroi

The cranial MDs were initially formed in *Gata2^cKO^*mesonephroi (Fig. 2A), and MD degeneration has been predominantly linked with AMH action. To rule out the involvement of AMH potentially produced from fetal ovaries or other organs in *Gata2^cKO^*female embryos, we performed ex vivo culture of E16.5 control and *Gata2^cKO^*mesonephroi without ovaries. After 5 days of culture, control MDs underwent elongation and formed an extensive coiled oviduct, whereas *Gata2^cKO^*MDs displayed a shortened oviduct with the loss of the cranial portion (Fig. 2B), recapitulating the in vivo phenotype (Fig. 1B). These observations indicate that cranial MD degeneration in *Gata2^cKO^*female embryos was not dependent on ovaries or other organs.

**Fig. 2.**
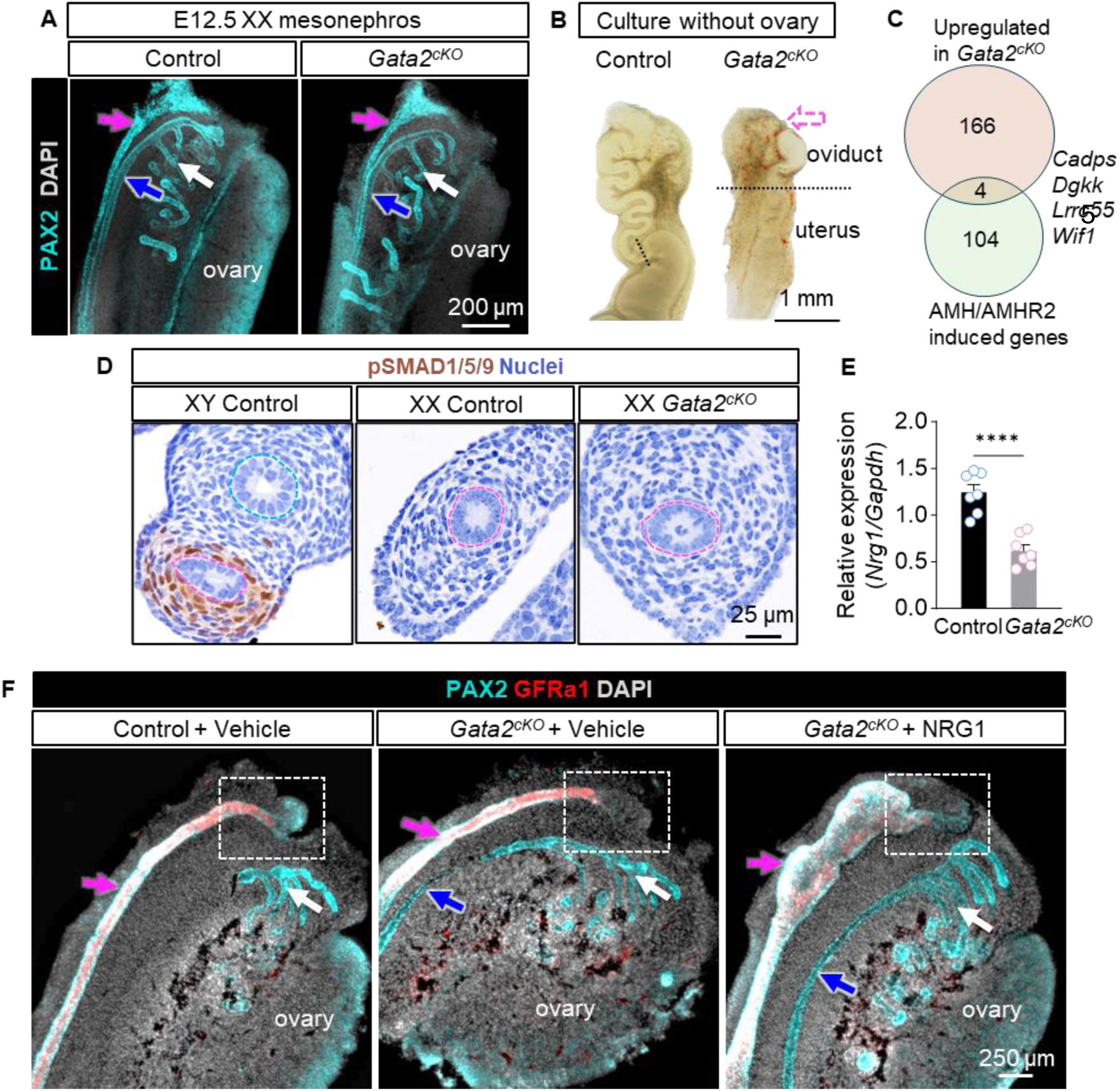
The loss of the cranial oviduct in *Gata2^cKO^* mice was not due to AMH/AMHR2 signaling but the reduced *Nrg1*. (**A**) Whole mount immunostaining of PAX2 in E12.5 XX control and *Gata2^cKO^* mesonephroi. Pink, blue, and white arrows: Müllerian duct (MD), Wolffian duct (WD), mesonephric tubules (MT), respectively. *n*=3 per group. (**B**) Bright field image of control and *Gata2^cKO^* mesonephroi without ovaries after 5-day culture. *n*=4 per group. Pink arrow: the position of the cranial oviduct lost in *Gata2^cKO^* embryos. Dashed line: the boundary between oviduct and uterus. (**C**) Venn diagram for gene overlap between upregulated DEGs in *Gata2^cKO^* cranial mesonephroi and AMH/AMHR2 induced genes. (**D**) Immunohistochemical staining of pSMAD1/5/9 in mesonephroi of E14.5 XY control, E16.5 XX control and *Gata2^cKO^*embryos, *n*= 3 per group. (**E**) Relative expression of *Nrg1* by RT-qPCR in XX control and *Gata2^cKO^* mesonephroi at E16.5, *n*=7 per group. \*\*\*\**P* <0.0001, Welch’s t test. (**F**) Whole-mount co-immunostaining of PAX2 and GFRa1 in E14.5 XX mesonephroi after 2 days of culture in the presence of vehicle or NRG1. *n*=3 per group. Pink, blue and white arrows: MD, WD, and MT, respectively. White dashed rectangle: the cranial region of MDs.

To determine any AMH signaling activities in cranial MDs of *Gata2^cKO^* female embryos, we performed bulk RNA-seq on cranial MDs from control and *Gata2^cKO^* embryos, yielded 170 upregulated genes in *Gata2^cKO^* (Data S1). Overlapping the list of these upregulated genes with AMH/AMHR2-induced genes (Data S2) (*9, 24*) resulted in only four genes (*Cadps*, *Dgkk*, *Lrrc55*, and *Wif1*) (Fig. 2C), none of which play a significant role in female reproductive tract development based on the lack of reproductive phenotypes in their individual knockout mice (*25–28*). Furthermore, we examined the expression of phosphorylated SMAD1/5/9 (pSMAD1/5/9), a key intracellular mediator of AMH signaling in MD regression (*29, 30*). In male control embryos with active AMH signaling, pSMAD1/5/9 was detected in the nuclei of MD mesenchyme. However, its expression was absent in MD mesenchyme in both control and *Gata2^cKO^* female mesonephroi, indicating that AMH signaling was not active in cranial MDs of *Gata2^cKO^*female embryos (Fig. 2D). Taken together, these results demonstrated that the cranial MD degeneration in the *Gata2^cKO^*female embryo is not due to ectopic AMH signaling.

### Contribution of reduced *Nrg1* to the *Gata2^cKO^* phenotype

Secreted ligands mediate the mesenchymal action in regulating the fate and differentiation of epithelial cells. Overlapping differentially expressed genes (DEGs) between cranial MDs from control and *Gata2^cKO^*mesonephroi (Data S1) with the mouse ligand atlas (*31*) yielded 13 secreted ligands that potentially involve in *Gata2^cKO^* MD degeneration (table S3). Based on the lack of phenotypes in reproductive organ development in their mutant mice, we deduced that 12 of them did not play any significant roles in MD development (table S3). We could not rule out the involvement of the downregulated gene *Nrg1* (Neuregulin 1) (Fig. 2E and table S3), a member of the epidermal growth factor (EGF) family of peptide growth factors. *Nrg1^-/-^* embryos died of heart malformation at E10.5-11.5, and conditional *Nrg1* knockout demonstrated its critical roles in fetal growth and morphogenesis of the testis and heart (*32, 33*). NRG1 signals through ErbB receptor tyrosine kinases to exert intracellular biological effects (*34*). The heterodimer of ErbB3 and ErbB2 that can mediate NRG1 actions is expressed in MD epithelium, as shown in prior studies (*32*) and GUDMAP (*35, 36*). Therefore, we were curious whether *Nrg1* downregulation could be the underlying cause of cranial MD regression in *Gata2^cKO^* embryos. To test this hypothesis, we performed ex vivo culture of E14.5 mesonephroi from control and *Gata2^cKO^*female embryos with vehicle or recombinant NRG1. After 2 days of culture, we performed whole-mount double immunostaining for epithelial marker PAX2 and MD epithelial marker GFRα1(GDNF family receptor α-1) (*37*) to visualize the development of the MDs (Fig. 2F). In vehicle-treated groups, PAX2 and GFRα1 expression were detected in the cranial MDs of control mesonephroi, while their expression was absent in the cranial region of *Gata2^cKO^* mesonephroi, indicating the degeneration of cranial MDs. In the NRG1-treated group, NRG1 supplementation promoted MD’s growth and lumen expansion in *Gata2^cKO^* mesonephroi, indicating MD’s responsiveness to NRG1 treatment. At the cranial region, NRG1 supplementation restored the expression of PAX2 and GFRα1, indicating the rescue of the *Gata2^cKO^* phenotype (Fig. 2F). Collectively, our results demonstrated that the reduced *Nrg1* contributed to the degeneration of cranial MDs in *Gata2^cKO^* female embryos.

### Defective uterine epithelial differentiation in *Gata2^cKO^* females

Unlike the oviductal phenotype, *Gata2^cKO^*uteri were maintained. To elucidate any impacts on uterine development in *Gata2^cKO^*females, we performed RNA-seq to examine transcriptomic differences between control and *Gata2^cKO^* caudal MDs at E16.5. Among the 537 DEGs, 60 (11.8%) were overlapped with 301 DEGs from cranial MDs (future oviduct) (fig. S5 and Data S1), suggesting regionally distinct transcriptomic changes upon *Gata2* deletion. The downregulated *Nrg1* that contributed to the cranial oviductal degeneration was not changed in the caudal MD, while *Gfra1,* the MD epithelial marker (*37*), was significantly downregulated only in the *Gata2^cKO^* caudal MD. Whole-mount immunostaining confirmed the lack of GFRa1 expression in the caudal MD (uterine region) (Fig. 3A), suggesting a possible loss of MD epithelium identity. Postnatal loss of *Gata2* in the whole uterus leads to luminal-to-basal epithelial transition (*38*). We therefore examine the expression of P63, the master regulatory transcriptional factor of the luminal-to-basal epithelial transition (*39–41*). Control and *Gata2^cKO^*oviducts did not display any P63 expression at PND0 and PND21 (fig. S7). P63 expression was not detected in the control uterus (Fig. 3B). However, in *Gata2^cKO^* uteri, P63 expression appeared in some epithelial cells at PND0 and became prevalent in the basal layer between the basement membrane and apical epithelia at PND21 (Fig. 3B). An important postnatal development in the uterus is glandular epithelial differentiation, which is driven by FOXA2 (*42–44*). *Gata2^cKO^* uteri at PND21 did not form uterine glands with FOXA2 expression limited to the apical epithelial on the mesometrial side (Fig. 3C). Collectively, these results demonstrated that mesenchymal *Gata2* deletion disrupts uterine epithelial differentiation.

**Fig. 3.**
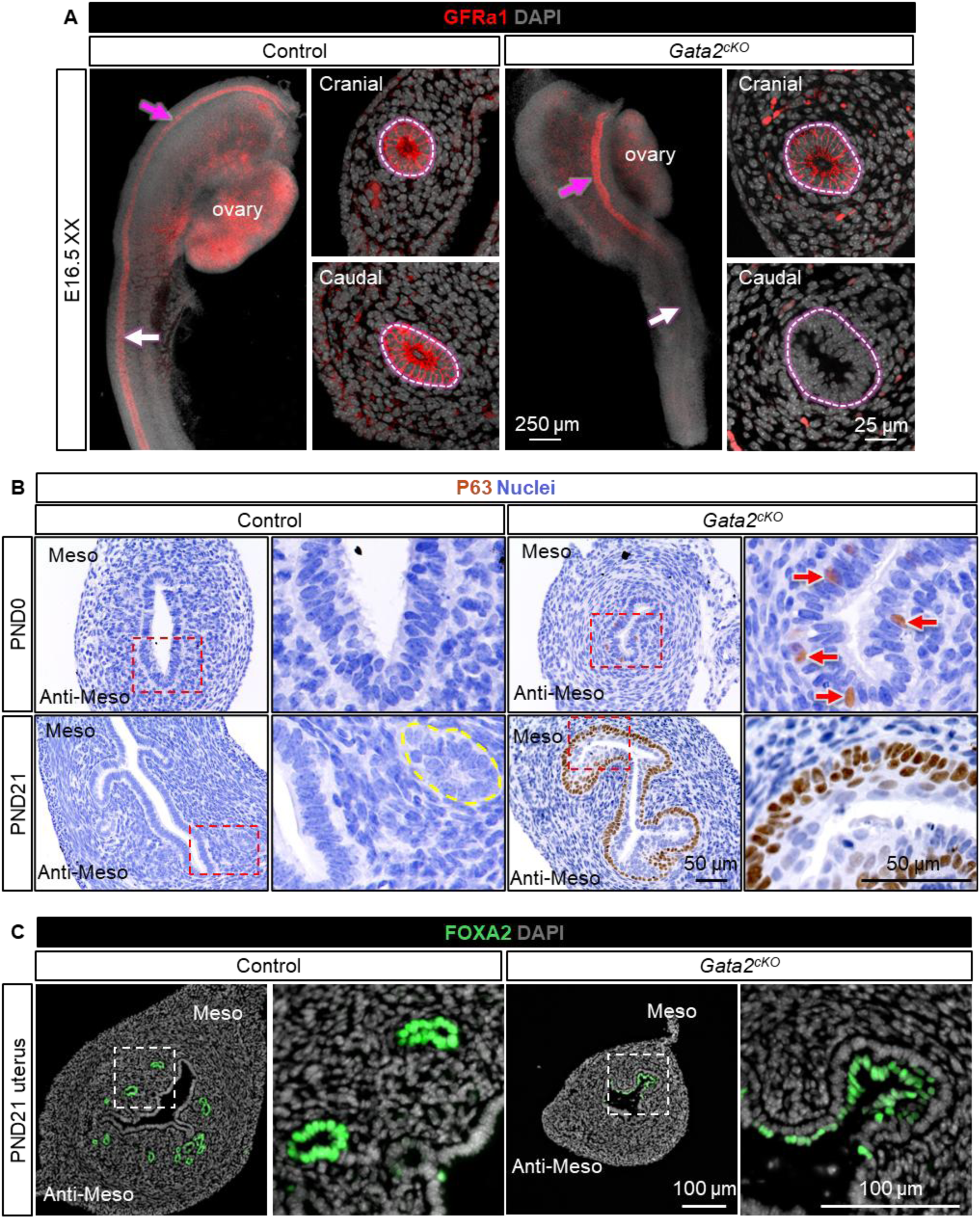
*Gata2^cKO^* driven by *Osr2-Cre* led to columnar-to-basal cells transition and defective gland formation in the uterus. (**A**) GFRa1 immunostaining on whole-mount tissue and cross-sections of XX control and *Gata2^cKO^*mesonephroi at E16.5. *n*=5 for control group, *n*=6 for *Gata2^cKO^*group. Pink and white arrows: cranial (oviductal) and caudal (uterine) region, respectively. (**B**) P63 immunohistochemical staining of control and *Gata2^cKO^* uteri at PND0 and PND21. The red dashed rectangle indicates the enlarged area. Yellow dashed line: uterine gland. *n*=3-4 per group. (**C**) FOXA2 immunofluorescent staining on the cross-sections of control and *Gata2^cKO^* uteri at PND21. *n*=3 per group. Meso: Mesometrial side; Anti-Meso: Anti-Mesometrial side.

### Impairment of uterine but not oviductal differentiation upon epithelial *Gata2* deletion

GATA2 expression appeared in uterine epithelia at E16.5 and PND0 (Fig. 1A), and *Osr2^Cre+^*lineage cells were observed in both the oviductal and uterine epithelia (fig. S2B). These observations prompted us to investigate any functional significance of epithelial *Gata2* and determine whether epithelial *Gata2* contributed to the oviductal and uterine phenotypes (Fig. 1B and Fig. 3, B and C) in our *Osr2-Cre:Gata2-flox* mouse model. We therefore generated an epithelium-specific *Gata2* knockout mouse model (*Pax8^Cre+^:Gata2^f/f^,* Epi-*Gata2^cKO^*) using *Pax8-Cre,* which targets the MD epithelium starting from E13.5 (*45–47*). At PND21, oviducts from Epi-Control (*Pax8^Cre+^:Gata2^f/+^*) and Epi-*Gata2^cKO^* females were morphologically and histologically indistinguishable with comparable lengths and distribution patterns of the secretory and ciliated oviductal epithelia in the infundibulum, ampulla and isthmus region (Fig. 4A and fig. S6, A and B). Epi-*Gata2^cKO^* oviducts did not exhibit P63 expression, like those of *Osr2-Cre*-driven *Gata2^cKO^* females (fig. S7). While PND21 uteri were morphologically comparable between Epi-Control and Epi-*Gata2^cKO^* females (fig. S8). Epi-*Gata2^cKO^* uteri exhibited some P63+ basal cells on the mesometrial side (Fig. 4B) and failed to develop uterine glands with the uterine gland marker FOXA2 ectopically expressed in the apical epithelia on the mesometrial side (Fig. 4C). Taken together, these results demonstrated that: (1) epithelial *Gata2* is dispensable for oviductal epithelial differentiation and does not contribute to the cranial MD degeneration; (2) epithelial *Gata2* regulates uterine epithelial differentiation.

**Fig. 4.**
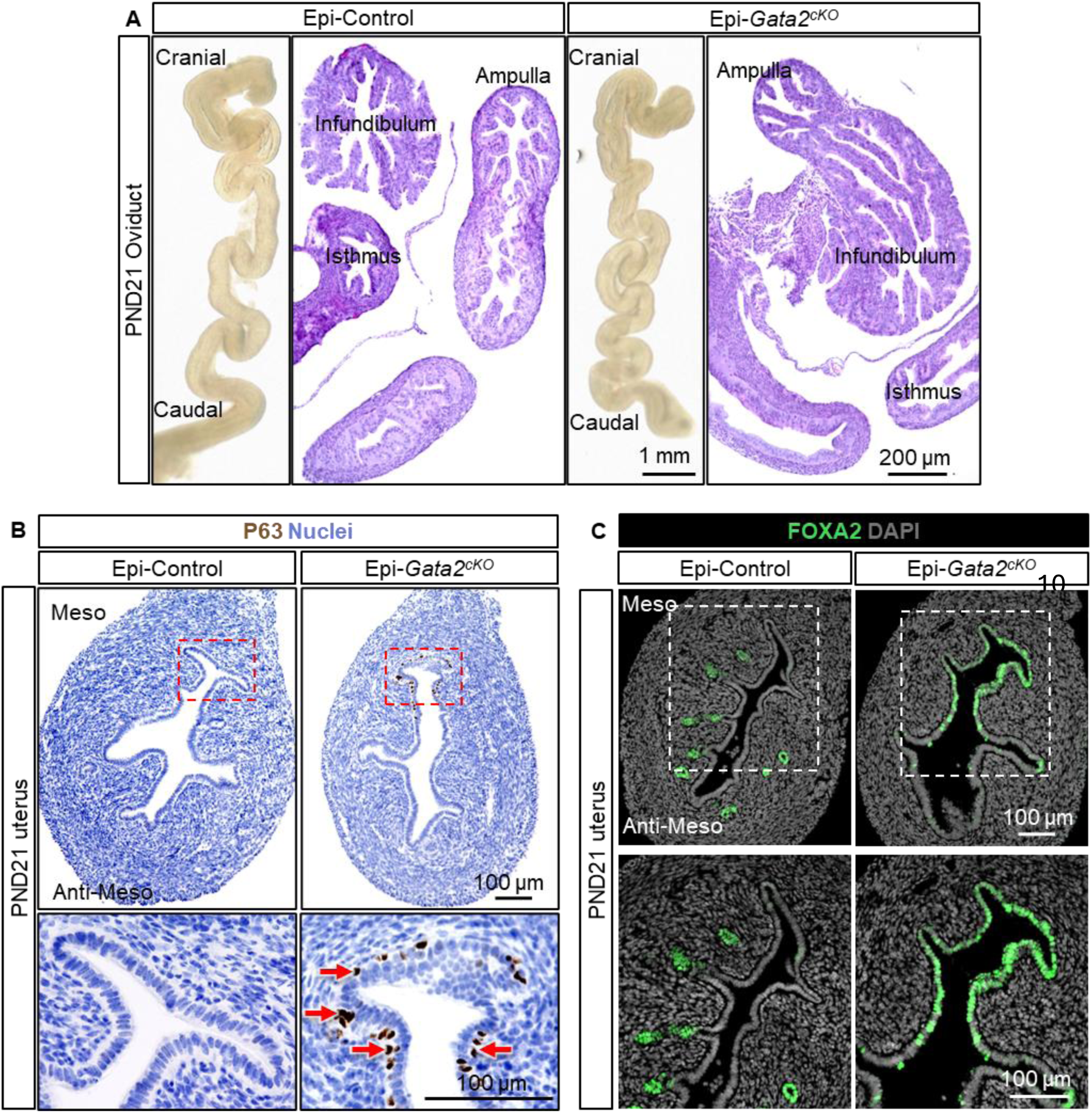
Deletion of epithelial *Gata2* using *Pax8-Cre* did not affect oviductal development but caused defective uterine epithelial differentiation. (**A**) Bright-field images and H&E staining of Epi-Control and Epi-*Gata2^cKO^* oviducts at PND21. (**B**) Immunohistochemical staining of P63 on the cross-sections of PND21 Epi-Control and Epi-*Gata2^cKO^* uteri. (**C**) Immunofluorescent staining of FOXA2 on the cross-sections of PND21 Epi-Control and Epi-*Gata2^cKO^* uteri. *n*=5 for Epi-Control, *n*=3 for Epi-*Gata2^cKO^*, for (A), (B) and (C). Meso: Mesometrial side; Anti-Meso: Anti-Mesometrial side.

## Discussion

Our study demonstrated region- and cell type-specific functions of *Gata2* during MD development. In the oviduct, mesenchymal *Gata2* promotes the maintenance of its cranial region while epithelial *Gata2* has a dispensable role in its development. In the uterus, both mesenchymal and epithelial *Gata2* safeguard the identity and differentiation of uterine luminal epithelium, preventing the transition from the columnar to basal epithelia.

### Mesenchymal *Gata2* promotes the maintenance of the cranial oviduct

We identify a previously unrecognized requirement for mesenchymal *Gata2* in MD maintenance, particularly within the cranial region. Mesenchymal *Gata2* is known to regulate epithelial growth and differentiation in several organs, including the WD (*48*), ureter (*13*), and inner ear (*14*). Loss of mesenchymal *Gata2* in these tissues inhibits ductal growth and/or impairs morphogenesis. Here, we showed that the loss of mesenchymal *Gata2* not only impaired MD growth but also led to selective degeneration of the cranial MD, a phenotype not observed caudally. It was previously reported that *Gata2* deletion in the urogenital system in compound *Gata2* mutant embryos led to defective WD and MD elongation during formation (*16*); however, the defective MD formation is probably secondary to impaired WD development which guides MD elongation (*49*). Therefore, our study reveals a unique, mesenchyme-specific role for GATA2 in MD maintenance following its formation.

The selective degeneration of the cranial MD implies the existence of a regionally distinct developmental program. Indeed, murine oviductal epithelial cells are derived from two distinct lineages: WT1+ in the cranial epithelia and PAX2+ in the caudal epithelia (*47*). Furthermore, the adult cranial oviduct is specifically enriched with *Slc1a3+* epithelial progenitor/stem cells (*50*). Beyond molecular markers, cranial and caudal oviducts are distinguished by their vasculature: the cranial region is supplied by the ovarian artery, whereas the caudal region depends on the uterine artery (*51*). Therefore, these findings highlight differences between the cranial and caudal oviducts in molecular regulatory programs and intrinsic properties.

The cranial MD regression in *Gata2^cKO^*female embryos occurs independently of AMH signaling. Our mechanistic analyses identified NRG1, a mitogenic growth factor, as a key downstream effector of mesenchymal GATA2. NRG1 has been suggested to mediate mesenchymal-epithelial crosstalk in postnatal oviductal development (*52*). Based on the ENCODE database, the GATA2 binding motif is found in the *Nrg1* promoter region, suggesting that *Gata2* may directly regulate *Nrg1* mRNA transcription (*53*). Both *GATA2* and *NRG1* are expressed in the mesenchyme of cranial oviducts in human embryos (*54–56*), raising the possibility of their potential roles in human MD maintenance and development.

Our findings support a model where mesenchymal GATA2 acts a gatekeeper, protecting the cranial MD from inappropriate degeneration in females. In male embryos, however, this program is overridden by testis-derived AMH. AMH-induced MD regression proceeds in a cranial-to-caudal direction in rats, correlating with a gradient of AMH type I and II receptors and the subsequent spreading of cellular apoptosis (*57, 58*). In addition, AMH reaches higher concentrations in the cranial region than in the caudal region because testis-derived AMH can infiltrate the mesonephros through the cranial region (*59, 60*), This cranial enrichment of AMH signaling likely outcompetes GATA2-mediated maintenance, thereby triggering the onset of cranial MD regression in males.

### Mesenchymal GATA2 safeguards uterine epithelial identity

Another important finding is that mesenchymal *Gata2* deletion leads to a loss of uterine epithelial identity. We first noted a marked reduction in the expression of the MD epithelial marker GFRa1, the co-receptor for GDNF (glial cell line-derived neurotrophic factor). Formation of the GFRa1-GDNF complex promotes the dimerization and activation of RET receptor tyrosine kinase, thereby inducing downstream intracellular signaling (*61*). *Gdnf* has been identified as a marker for caudal MD epithelium during MD elongation in humans (*56*), but *Ret* and *Gdnf* expression in MD or its mesenchyme remains undefined. In the kidney, GDNF/ GFRa1/RET signaling drive ureteric bud outgrowth from the WD (*62*); however, their functions in MD development are largely undefined.

The columnar-to-basal epithelial transition in our *Gata2^cKO^* model aligns with findings from a prior study in which *Pgr-Cre* was used to delete *Gata2* in the postnatal uterus (*38*). It is noteworthy to point out the differences and similarities between the two models. *Osr2-Cre* activity begins as early as E12.5, whereas *Pgr-Cre* activity is initiated postnatally around PND14 (*63*). Reflecting these temporal differences in Cre recombinase activation, P63+ basal-like cells emerge earlier in our model compared with *Pgr-Cre-*mediated *Gata2* knockout. Despite the distinct timing of *Gata2* deletion, both models exhibit a columnar-to-basal epithelial transition, suggesting the conserved requirement for *Gata2* in maintaining uterine columnar epithelial identity across both fetal and postnatal stages.

The molecular mechanism by which mesenchymal GATA2 prevents aberrant basal cell differentiation in the uterus remains to be further investigated. Under normal conditions, P63+ basal cells and epithelial stratification are restricted to the vagina (*64*). Activin A (*Inhba*), *Fgf7/10* or *Bmp4* are known to regulate P63 induction and basal cell differentiation in vaginal epithelium. However, none of these factors emerged differentially expressed in our E16.5 *Gata2^cKO^*transcriptomic analysis, suggesting that the mechanisms driving basal cell formation in the *Gata2^cKO^* uterus are distinct from those governing normal vaginal epithelial differentiation. Vitamin A (retinoic acid signaling) deficiency is another known cause of columnar-to-basal epithelial transition in the mouse uterus (*65, 66*). However, the expression of retinoic acid downstream targets, including *Hoxa4*, *Hoxb5*, *Hoxc4*, and *Hoxc5* (*67*), was comparable between control and *Gata2^cKO^* uteri, indicating that RA signaling was unperturbed. Ectopic expression of *Hoxa13* in place of *Hoxa11* in the uterus (*68*), or concurrent deletion of *Hoxa9, 10,11* (*69*) can induce transformation of uterine epithelial cells to basal cells. Yet, *Hoxa9, 10 or 13* expression levels were not altered in *Gata2^cKO^* uteri. The only differentially expressed *Hox* gene was *Hoxa11*. It has been documented that *Hoxa11* mutant adult mice lack uterine gland formation (*70*), suggesting that reduced *Hoxa11* expression may contribute to defective uterine gland formation observed in *Gata2^cKO^*uteri.

### Critical roles of epithelial GATA2 in uterine not oviductal differentiation

In contrast to the dramatic phenotype resulting from mesenchymal *Gata2* deletion, Epi-*Gata2^cKO^* females develop morphologically normal oviducts with intact overall tissue architecture. Our analyses revealed that GATA2 is not expressed in the fetal oviductal epithelium; in adults, *Gata2/GATA2* expression seems to be enriched in the oviductal stroma, as supported by published mouse (*71*) and human datasets (*72*). These observations suggest that the major GATA2 action required to maintain the cranial MD originates from the mesenchyme rather than epithelium, consistent with the established paradigm that mesenchymal cues direct epithelial differentiation and morphogenesis (*73*).

The uterine phenotypes of Epi-*Gata2^cKO^*females are considerably less severe than those of *Osr2-Cre*-driven *Gata2^cKO^*. For instance, uterine size, especially the mesenchymal compartment, remains substantially larger in Epi-*Gata2^cKO^* uteri, suggesting that the mesenchymal proliferation is primarily dependent upon mesenchymal *Gata2* rather than epithelial *Gata2-*driven paracrine signaling. Moreover, Epi-*Gata2^cKO^*uteri exhibit only sparse P63+ cells beneath the luminal epithelium on the mesometrial side. This contrasts with the extensive and multilayered basal cell population in the absence of mesenchymal *Gata2*. These observations underscore the dominant role of mesenchymal *Gata2* in restraining columnar-to-basal cell transition.

Uterine gland development is disrupted in both *Gata2^cKO^*and Epi-*Gata2^cKO^* models. Although the glandular epithelial marker FOXA2 is present in the epithelia on the mesometrial side in both mutants, glandular morphogenesis does not occur. These observations suggest that while glandular epithelial specification may initiate, subsequent morphogenetic steps such as budding, elongation, and branching fail to proceed (*74*). Notably, both GATA factors and FOXA2 have been implicated in transcriptional programs associated with glandular epithelial differentiation (*75*), raising the possibility of functional interplay between these two transcription factors during epithelial cell lineage progression.

The clustering of FOXA2+ cells specifically on the mesometrial side is intriguing. In rodents, uterine glands typically develop in the lateral and anti-mesometrial regions, whereas the mesometrial area is largely devoid of glands (*44*). Spatially restricted WNT signaling has been proposed to confine gland formation to appropriate regions. Therefore, the abnormal localization of FOXA2+ epithelial cells may reflect disrupted WNT-mediated spatial cues in the absence of either epithelial or mesenchymal *Gata2*.

Congenital anomalies of MDs represent a major cause of female infertility (*76*), affecting approximately 0.1–3.0% of live births (*77*) and costing over $40 billion annually worldwide (*78*). These statistics highlight the urgent need to better understand the mechanisms governing normal female reproductive tract development. Our findings challenge the traditional view that MD maintenance in females is a passive outcome that occurs in the absence of AMH signaling. We demonstrate that cranial MD maintenance in females is an active process promoted by mesenchymal GATA2, which acts at least in part through induction of *Nrg1* signaling. Furthermore, we uncover the uterus-specific function of GATA2 in preventing columnar-to-basal epithelial transformation. Taken together, our work provides new insights into the molecular regulation of female reproductive tract development and potentially advances our understanding of the pathogenesis of Müllerian anomalies in humans.

## Acknowledgments

We would also like to thank Dr. Rulang Jiang from Cincinnati Children’s Hospital Medical Center, Division of Developmental Biology for providing *Osr2-Cre* mice; Mutant Mouse Regional Resource Center for providing *Gata2-flox* mice; High Throughput Computing Center, UW-Madison for providing the sequencing analysis platform.

## Author contributions

Conceptualization: SJ, FZ

Methodology: SJ, FZ

Investigation: SJ

Visualization: SJ, AF

Funding acquisition: FZ

Project administration: SJ

Supervision: FZ

Writing – original draft: SJ, FZ

Writing – review & editing: SJ, AF, WYB, FZ

## Data, code, and materials availability

All data is available in the main text and the supplementary materials. The RNA-seq data was deposited in the GEO under the accession number GSE324404.

## Supplementary Materials

### Materials and Methods

#### Mice

The *Osr2-Cre* (knock-in) strain was provided by Dr. Rulang Jiang at the Cincinnati Children’s Hospital Medical Center and maintained in its original genetic background (129/Sv × C57BL/6J) (*21*). The *Gata2-flox* strain was purchased from the Mutant Mouse Resource and Research Center of the University of Missouri (MMRRC:030290). *Pax8-Cre* (028196) and *Rosa-tdTomato* (007909) strains on the C57BL/6J genetic background were purchased from Jackson Laboratories (Bar Harbor, ME). Timed mating was set up by housing stud males with 2-3 sexually mature females (2 to 6 months old) in the late afternoon and the morning of plug detection was designated as embryonic day 0.5 (E 0.5). Genotyping PCR was performed using primers listed in table S1. The following thermal cycle with Platinum II Taq Hot-Start DNA Polymerase (Invitrogen, 14966001) was used for genotyping: 94 °C for 2 min, followed by 34 cycles of [94 °C for 15 sec, 60 °C for 5 sec, and 68 °C for 15 sec], then 68 °C for 5 min. *Osr2^Cre+^*:*Gata2^f/+^*and *Osr2^Cre+^*:*Gata2^f/f^* were designated as control and *Gata2^cKO^*, respectively. *Pax8^Cre+^*:*Gata2^f/+^*and *Pax8^Cre+^*:*Gata2^f/f^* were designated as Epi-Control and Epi-*Gata2^cKO^*, respectively. All mouse procedures were approved by the University of Wisconsin-Madison (UW-Madison) Animal Care and Use Committees and followed UW-Madison approved animal study proposals and public laws.

#### Tissue processing

Tissues were fixed in 10% neutral buffered formalin (Leica, 3800598) overnight at room temperature, then processed and embedded as previously described (*79*). Paraffin sections were cut at 5 μm for staining.

#### Histology

H&E (Hematoxylin and eosin) staining was performed on paraffin sections according to an established protocol (*80*). In brief, sections were deparaffinized in xylene, rehydrated through a graded ethanol series, and stained with Harris hematoxylin (Sigma, HHS128) for 1 min. Following a brief rinse in running tap water, differentiation was carried out in acid ethanol and sections were washed in tap water. Counterstaining was performed with eosin Y (Leica Biosystems, 3801615) for 2 min. Sections were then dehydrated through graded ethanol, cleared in xylene, and mounted with Permount mounting medium (Fisher Chemical, SP15-500). Bright-field images were acquired using a Keyence BZ-X710 fluorescence microscope (Keyence, Osaka, Japan).

#### Immunofluorescence

Immunofluorescence staining was performed as previously described (*79*). Briefly, paraffin sections were deparaffinized, rehydrated, and then subjected to antigen retrieval (VECTOR, H-3300) using a microwave oven. After washing with PBST (1× PBS with 0.1% Triton X-100), sections were incubated in the blocking buffer (5% normal donkey serum in PBST) for 1 hour and then with a primary antibody in blocking buffer overnight at 4 °C. The next day, sections were washed three times with PBST, incubated with secondary antibodies for 1 hour at room temperature, and counterstained with DAPI (Thermo Fisher, 62248, 1:1000). Images were taken under a Leica TCS SP8 confocal microscope or a Leica Thunder Imaging System. The primary and secondary antibodies used were listed in table S2. The percentage of Ki67+ and cleaved-PARP1+ cells among the total number of cells (nuclei stained by DAPI) in the epithelial and mesenchyme compartments were quantified from three to six sections per mouse in each group using ImageJ (*79*).

#### Immunohistochemistry

Paraffin sections were deparaffinized, rehydrated, and subjected to antigen retrieval as described in the immunofluorescent staining section. Endogenous peroxidase was inhibited using 3% H_2_O_2_ (VWR, 470301-282). After washing with PBST, slides were incubated in the blocking buffer for 1 hour, then with primary antibody in blocking buffer overnight at 4 °C. After washing with PBST, sections were incubated with the secondary antibody for 30 minutes at room temperature, washed with PBST and then incubated with ABComplex/HRP (VECTOR, PK-6100) for 30 minutes at room temperature. Slides were treated with DAB (VECTOR, SK4100) for signaling development and counterstained with hematoxylin (Sigma, HHS128) for imaging under a Keyence microscope (BZ-X710). The primary and secondary antibodies used were listed in table S2.

#### Whole-mount immunostaining

Whole-mount immunostaining was performed as previously described (*81*). Briefly, either freshly dissected or cultured intact mesonephroi (with gonads attached) were fixed in 4% paraformaldehyde (PFA) overnight at 4 °C, then washed three times in 1× PBS and incubated with blocking solution for 1 hour on a platform rocker (Corning, 6703) at 50-60 RPM. The tissue was then incubated with primary antibody in blocking buffer overnight at 4 °C with rocking, followed by three washes with PBST and incubation with secondary antibody for 2 hours at room temperature with rocking. Whole-mount tissues were counterstained with DAPI (1:1000, Thermo Fisher, 62248) and imaged under a Leica TCS SP8 confocal microscope or a Leica Thunder Imaging System.

#### Real-time qPCR

The cranial and caudal regions of mesonephroi from E16.5 control and *Gata2^cKO^* embryos were separated at the flexura medialis (*82, 83*) and snap frozen on dry ice. RNA extraction was performed using the PicoPure RNA isolation kit (Life Technologies, KIT0204) according to the manufacturer’s protocol. The RNA concentration and purity were determined using a Nanodrop Spectrophotometer ND-1000, with both 260/280 and 260/230 ratios above 2.0. Complementary DNA (cDNA) was synthesized from 250 ng RNA per sample using the RT HT First Strand Kit (Qiagen, 330411) and used in a qPCR reaction with SYBR Green and specific primers (table S1). Reactions were run on a Bio-Rad CFX96 with the following program: initial denaturation at 95 °C for 2 min; 40 cycles of 95 °C for 5 s and 60 °C for 10 s (plate read); followed by melt curve analysis from 65 °C to 95 °C (0.5 °C increments every 5 s, plate read). Data were exported using Bio-Rad CFX Maestro 2.3 (v5.3.022.1030). All samples were duplicated and normalized to *Gapdh*. The relative gene expression was calculated using the 2^−△△Ct^ method.

#### Ex vivo organ culture

Our previously established protocol was used (*81*). Mesonephroi with or without attached ovaries were cultured on 0.4 µm Millicell-CM inserts (Millipore, PICM03050) in 6-well plates at 37 °C with 5% CO₂/95% air. The following culture media were used: 1 mL phenol red-free DMEM/F12 (Gibco, 21041025) supplemented with 100 U/mL penicillin-streptomycin (Gibco, 15070063) and 1% Insulin-Transferrin-Selenium (Gibco, 41400045). For NRG1 (Bio-Techne, 396-HB,100 ng/mL) treatment, sterile PBS with 1% BSA (Dot Scientific, DSA30075-25) was used as the vehicle. After culture, tissues were imaged under a Leica S9 stereo microscope and fixed in 4% PFA for subsequent whole-mount immunostaining.

#### RNA-sequencing and data analysis

Extracted RNA from E16.5 control and *Gata2^cKO^*cranial (oviduct region) and caudal (uterus region) mesonephroi were analyzed and sequenced (paired end, 150 nt) by Novogene. The RNA integrity number (RIN) in all samples was above 7.5. Raw read quality was assessed using fastQC (v0.12.1). After filtering low-quality, high-N, and short reads with fastp (v0.23.4) under default settings, reads were aligned to the mouse mm39 genome using STAR (v2.7.11b) and gene counts were generated using featureCounts (v2.0.6) with default parameters. Differential expression gene analysis was performed with DESeq2 (v1.42.0) under R program (v4.3.2) with the cutoff |log₂(FoldChange)| > 1 and padj < 0.05 (*Gata2^cKO^* vs. Control).

#### Statistical analyses

At least three biological replicates per group were used in each experiment and sample sizes are indicated in figure legends. Statistical analyses were performed with GraphPad Prism (version 10.6.1). Statistical significance between control and knockout groups was assessed by first applying a Shapiro-Wilk test for normality. If the data were normally distributed, a two-tailed unpaired Welch’s t test was used; if the data were nonparametric, a Mann-Whitney U test was performed. Results were shown as Mean ± SEM. The statistical significance was set at *P* < 0.05. The exact *P* values were indicated in the corresponding figure legends.

**Fig. S1.**
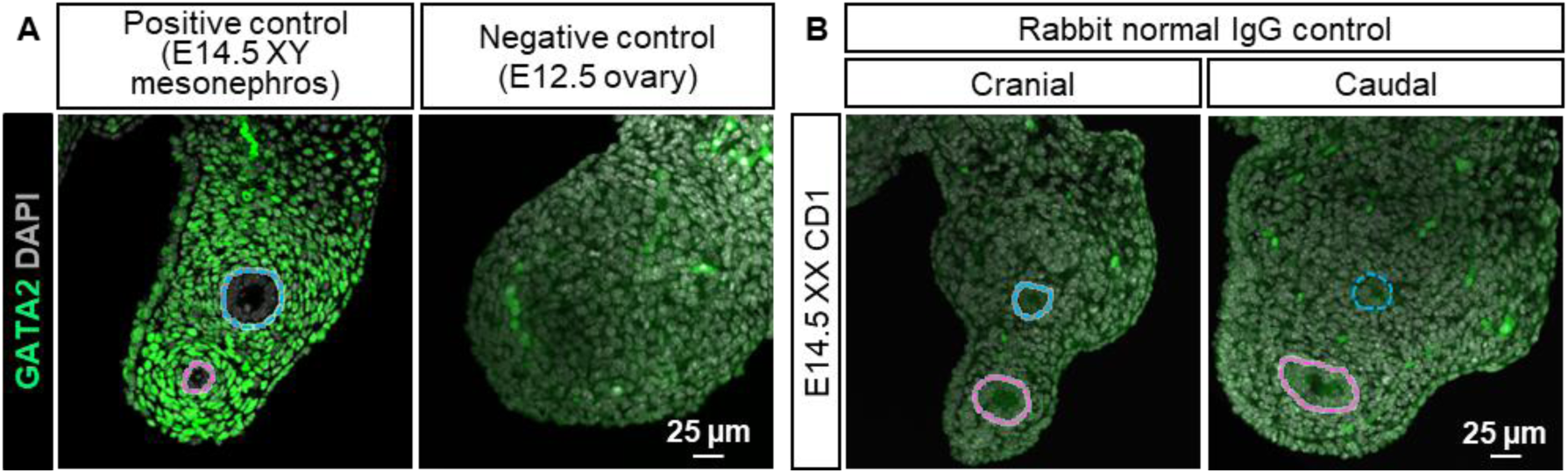
Controls for GATA2 immunostaining. (**A**) GATA2 immunofluorescence staining in E14.5 XY mesonephroi (*n*=6), and E12.5 ovary (*n*=2). (**B**) Immunofluorescence staining of normal rabbit IgG in E14.5 XX cranial and caudal mesonephroi. *n*=3 per region.

**Fig. S2.**
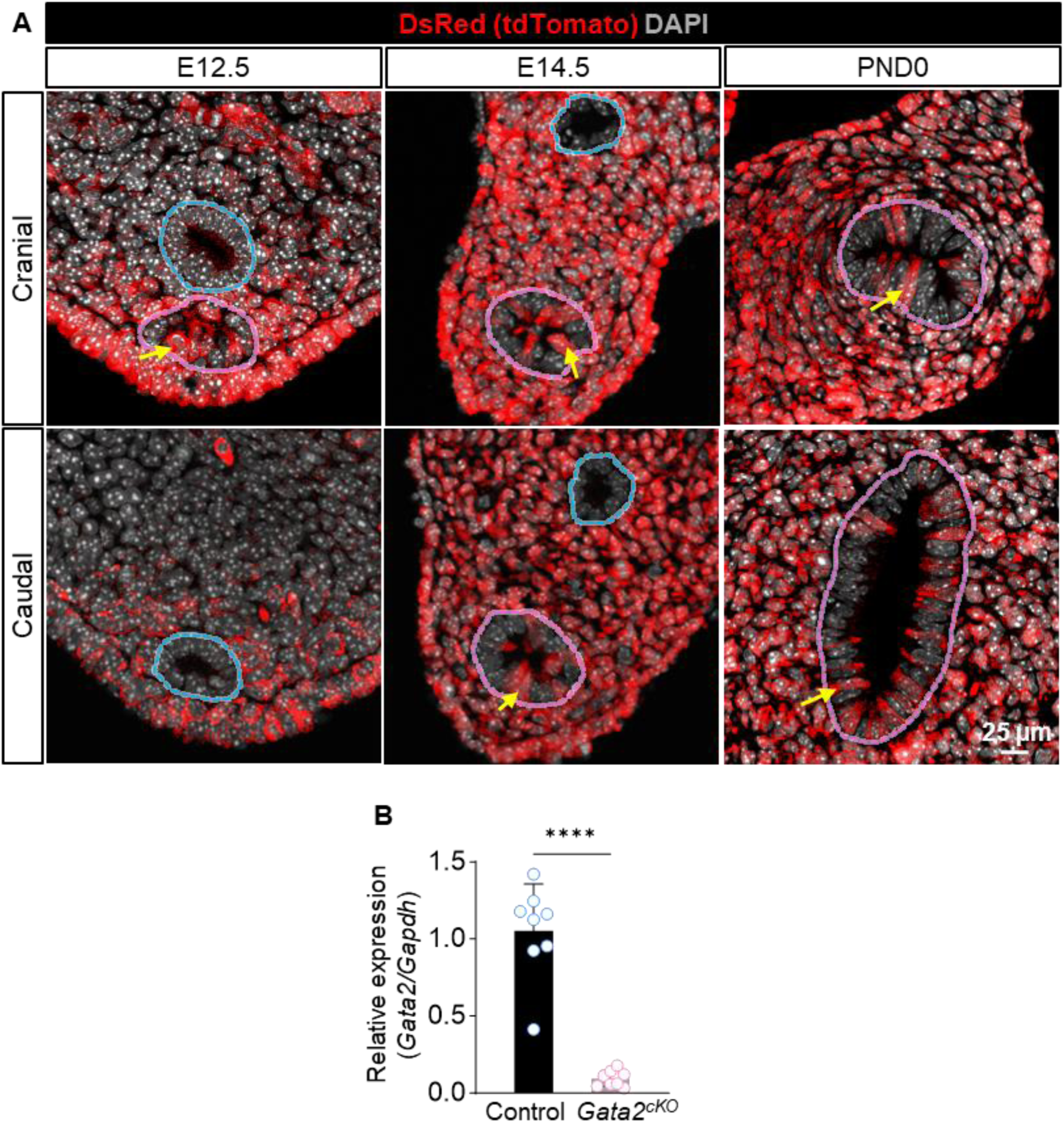
Characterization of *Osr2-Cre* activity. (**A**) Immunofluorescent staining of DsRed (tdTomato) on cross-sections of cranial and caudal mesonephroi at E12.5, E14.5 and PND0 *Osr2^Cre^*^+^:*tdTomato*^+^ female embryos. *n*=3 per timepoint. Blue and pink dashed lines: Wolffian and Müllerian duct, respectively. Yellow arrows indicate DsRed+ epithelial cells. (**B**) Relative expression of *Gata2* by RT-qPCR in E16.5 XX control and *Gata2^cKO^* cranial mesonephroi. *n*=8 per group. \*\*\*\**P*<0.0001, Welch’s t test.

**Fig. S3.**
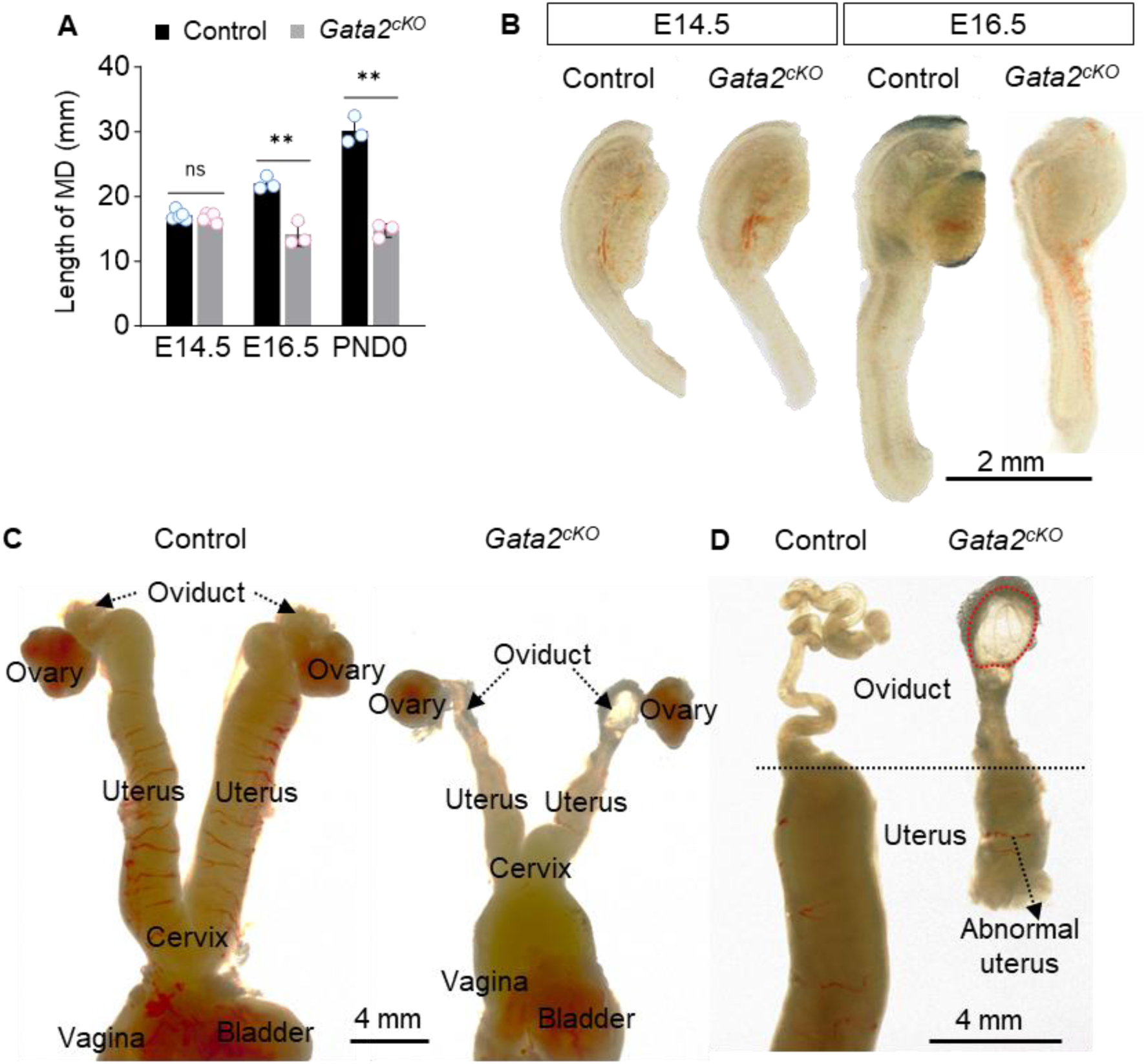
*Gata2^cKO^* driven by *Osr2-Cre* causes decreased Müllerian duct length and hydrosalpinx at adulthood. (**A**) Length of female mice Müllerian duct at E14.5 (*n*=5 per group, ns: not significant, *P*=0.4419, Welch’s t test), E16.5 (*n*=3 per group, \*\**P*=0.0065, Welch’s t test), and PND0 (*n*=3 per group, \*\**P*=0.0014, Welch’s t test). (**B**) Bright-field images of control and *Gata2^cKO^* female mesonephroi at E14.5 (*n*=5 per group), E16.5 (*n*=3 per group). (**C**) Bright-field images of control and *Gata2^cKO^* female whole reproductive tract in adult stage. The corresponding organ regions were indicated by the labels: ovary, oviduct, uterus, cervix, vagina and bladder. (**D**) Bright-field images of control and *Gata2^cKO^* female upper reproductive tract. Red dashed lines: fluid-filled oviduct (Hydrosalpinx), black dashed line: the boundary between oviduct and uterus. *n*=4 per group.

**Fig. S4.**
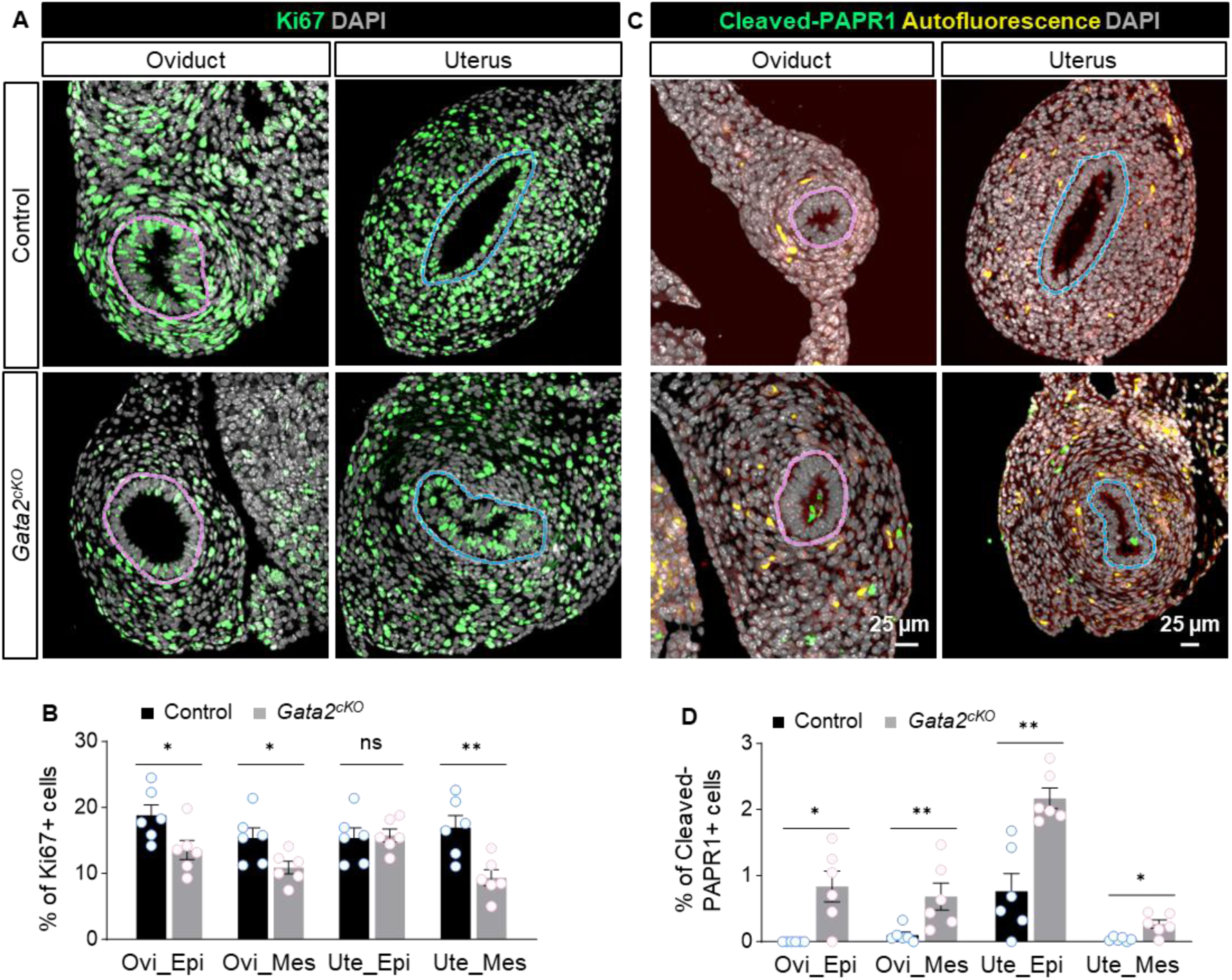
*Gata2^cKO^* driven by *Osr2-Cre* decreases proliferation and increases apoptosis in the epithelium and mesenchyme of Müllerian ducts. (**A & C**) Immunofluorescent staining of the proliferation marker Ki67 (**A**) and apoptotic marker cleaved-PARP1 (**C**) in the oviduct and uterus of control and *Gata2^cKO^*embryos at PND0. Pink and blue dashed lines: Müllerian and Wolffian ducts. (**B**) Percentage of Ki67+ cells in Ovi_Epi (\**P*=0.0352), Ovi_Mes (\**P* =0.0368), Ute_Epi (ns *P* =0.2772) and Ute_Mes (\*\**P* =0.0771) of control and *Gata2^cKO^* mice at PND0. Welch’s t test was used in each comparison. (**D**) Percentage of Cleaved-PAPR1+ cells in Ovi_Epi (\**P* =0.0152), Ovi_Mes (\*\**P* =0.0087), Ute_Epi (\*\**P* =0.0019) and Ute_Mes (\**P* =0.0169) of control and *Gata2^cKO^* mice at PND0. Mann-Whitney U test was used in Ovi_Epi and Ovi_Mes, Welch’s t test was used in Ute_Epi and Ute_Mes. *n*=6 per group in both (B) & (D). ns: nonsignificant. Ovi_Epi: oviductal epithelium, Ovi_Mes: oviductal mesenchyme, Ute_Epi: uterine epithelium, Ute_Mes: uterine mesenchyme.

**Fig. S5.**
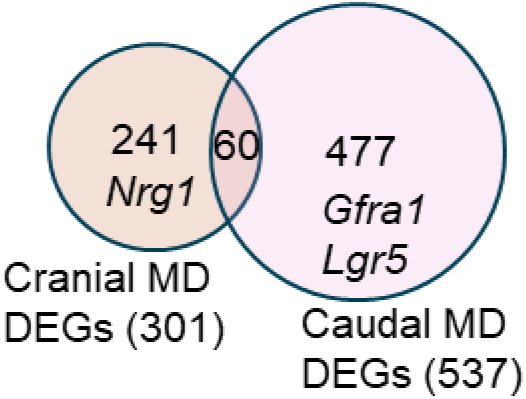
*Gata2^cKO^* driven by *Osr2-Cre* caused regional-specific DEGs. Venn diagram illustrates the overlapping between DEGs in *Gata2^cKO^* cranial and *Gata2^cKO^* caudal mesonephroi.

**Fig. S6.**
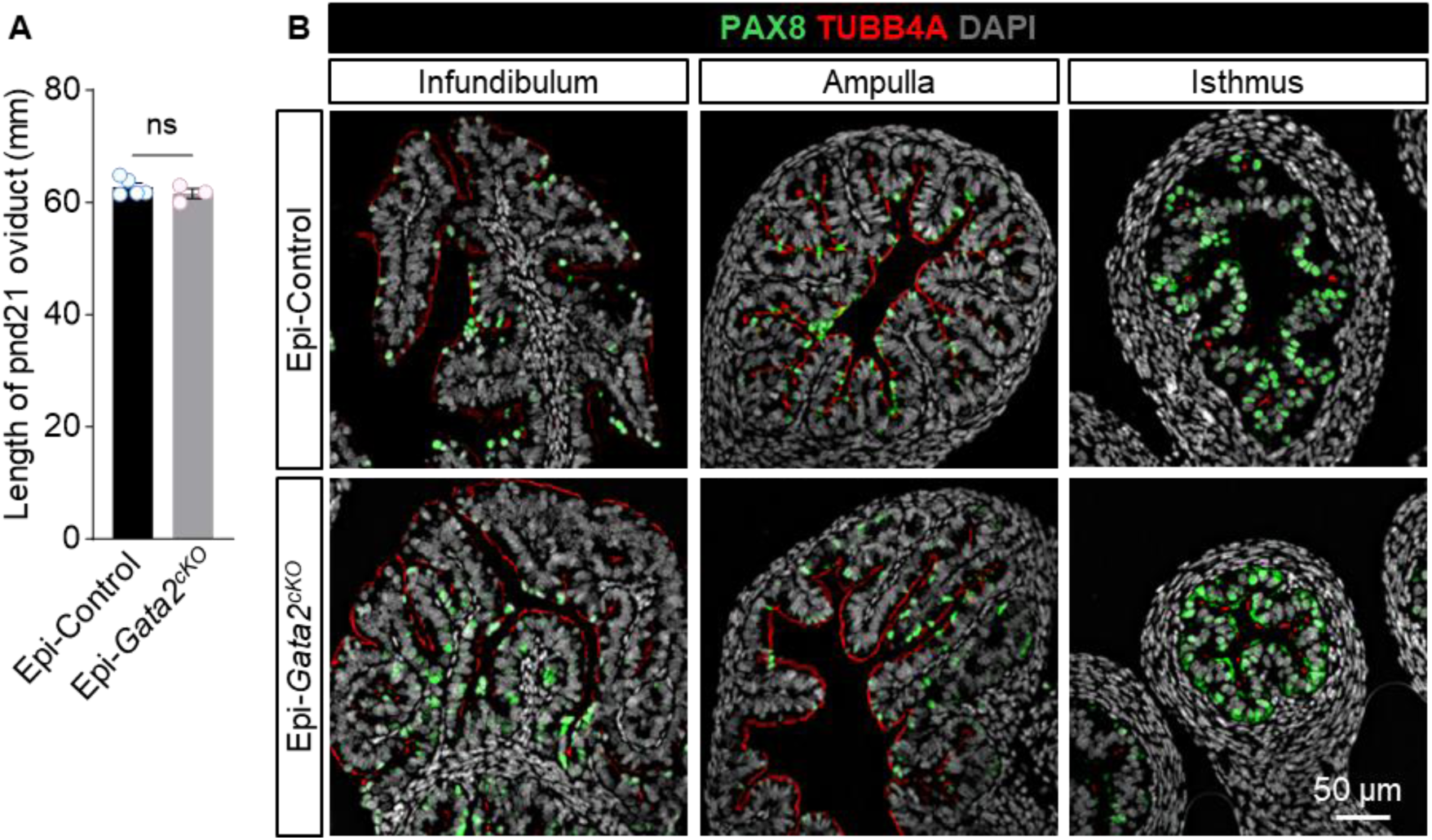
Deletion of epithelial *Gata2* using *Pax8-Cre* did not affect oviductal length or epithelial cell differentiation. (**A**) Length of Epi-Control (*n*=5) and Epi-*Gata2^cKO^*(*n*=3) oviducts at PND21. ns: not significant, *P*=0.3584, Welch’s t test. (**B**) Immunofluorescent staining of PAX8 (secretory cell marker) and TUBB4A (ciliated cell marker) in the infundibulum, ampulla and isthmus of PND21 Epi-Control and Epi-*Gata2^cKO^* oviducts. *n*=5 per group.

**Fig. S7.**
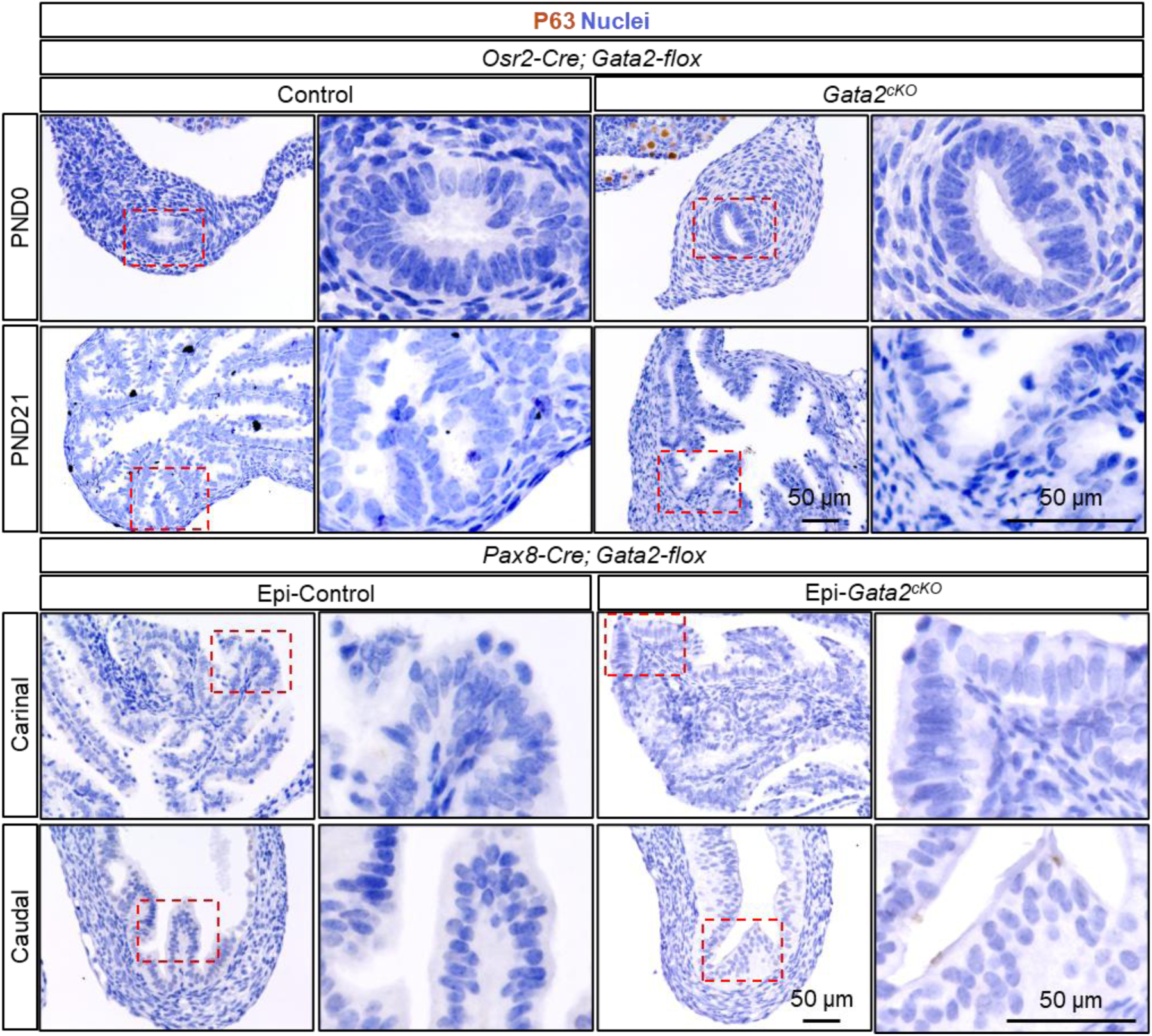
*Gata2* deletion in mesenchymal or epithelial cells did not cause columnar-to-basal cell transition in the oviduct. Immunohistochemistry staining of P63 in control and *Gata2^cKO^* oviducts at PND0 and PND21, and in Epi-Control and Epi-*Gata2^cKO^* oviducts at PND21. Red dashed line rectangle: the enlarged area. *n*=3-5 per group in each timepoint.

**Fig. S8.**
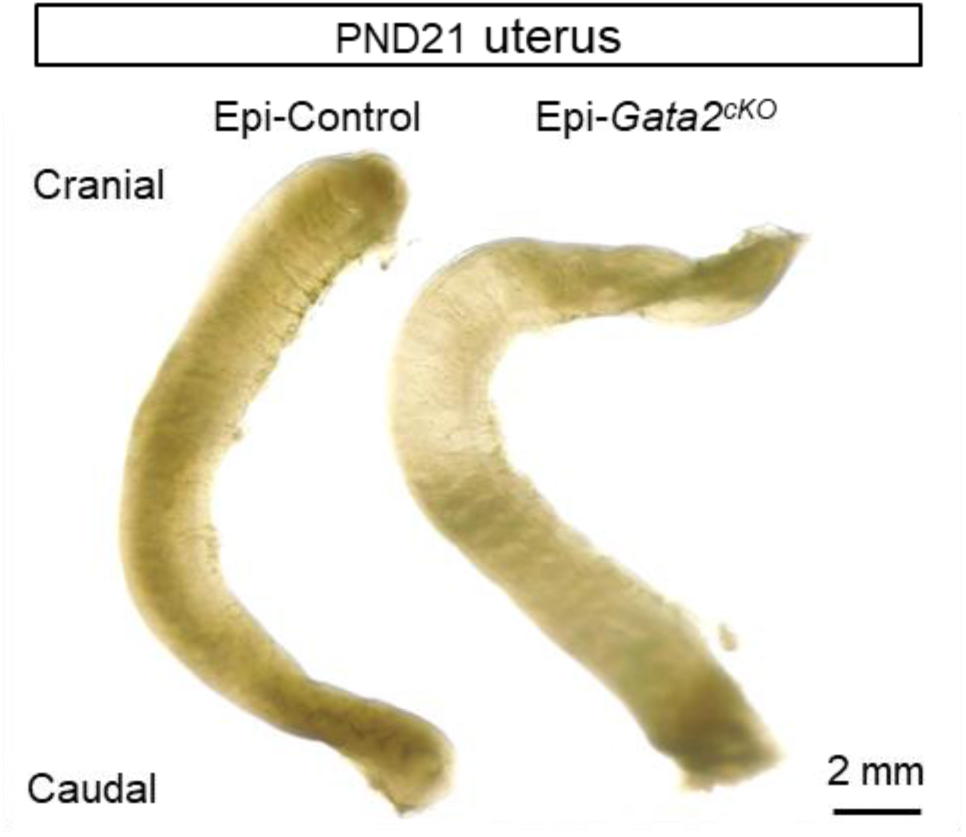
Epithelial *Gata2* knockout did not affect gross morphology of the uterus. Bright-field images of Epi-Control and Epi-*Gata2^cKO^* uteri at PND21. *n* =3 per group.

**Table S1.**
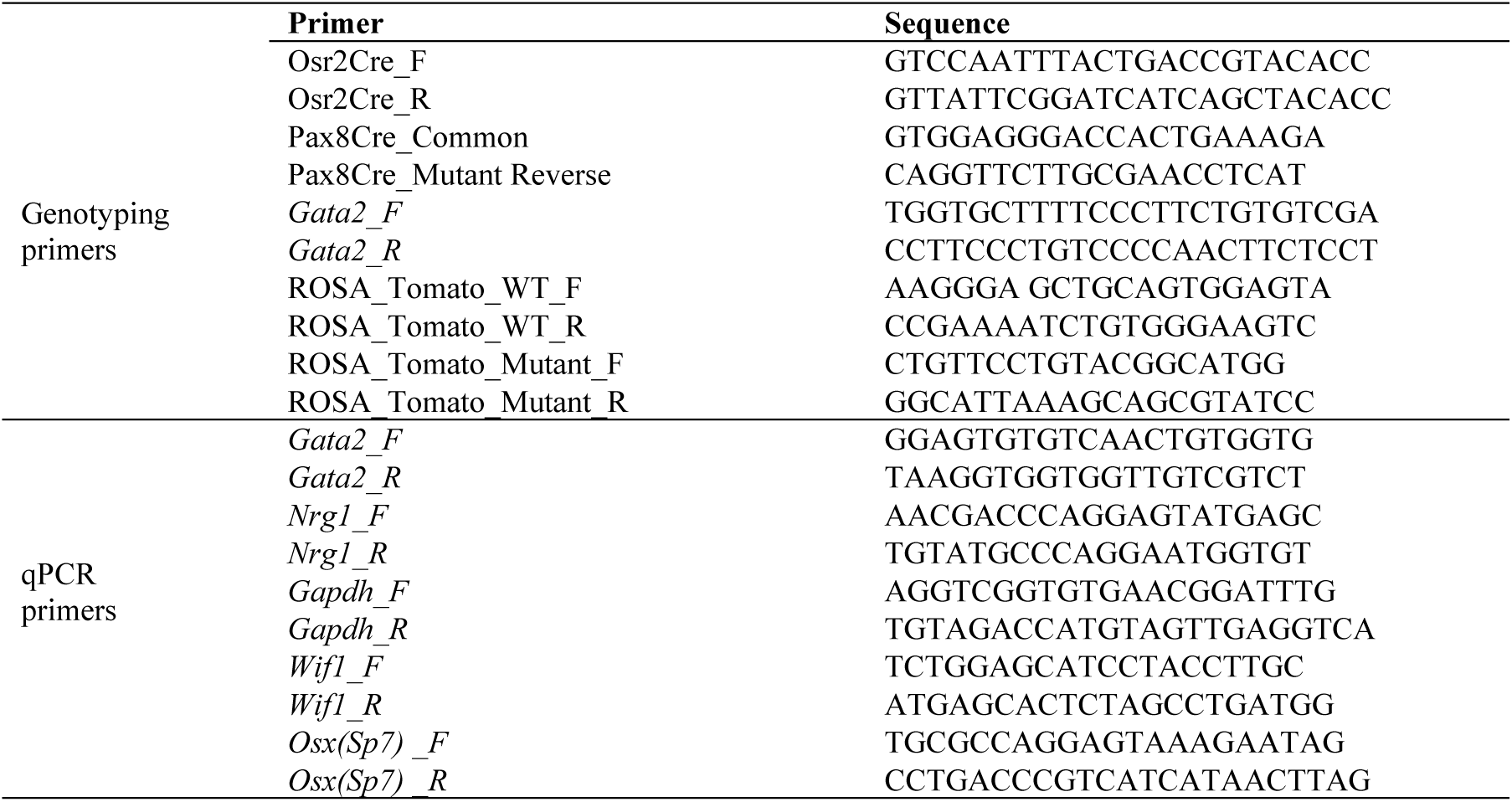
Primers used for genotyping and qPCR.

**Table S2.**
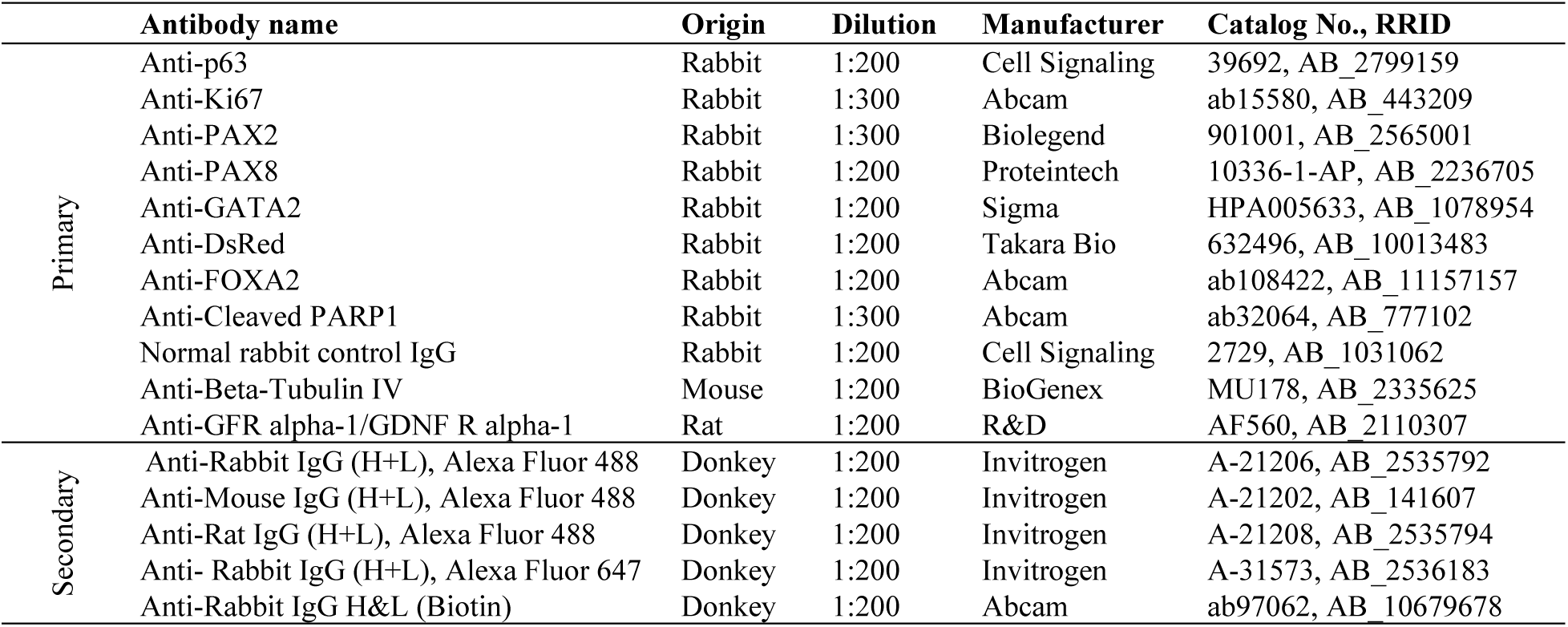
Antibody list.

**Table S3.** Phenotypes of 13 enriched Up- and Down-regulated ligands in *Gata2^cKO^*cranial Müllerian ducts.

|  | **Gene** | **Phenotype after mutation** | **Reference** |
| --- | --- | --- | --- |
| Down-regulated<br>in *Gata2<sup>cko</sup>* | *Wnt2* | Fertile (survivors); ~50% died perinatally | PMID: 8951051 |
|  | *Sfrp5* | Fertile, no developmental abnormalities | PMID: 17133501 |
|  | *Nrg1* | Embryonic lethality at E10.5 | PMID: 11395002 |
|  | *Bmp7* | Fertile (subfertile) | PMID: 28324064 |
|  | *Wnt6* | Fertile | PMID: 23175771 |
|  | *Nrtn* | Fertile | PMID: 10069332 |
|  | *Lrp1b* | Fertile | PMID: 15082773 |
| Up-regulated<br>in *Gata2<sup>cko</sup>* | *Ccn3(Nov)* | Fertile (survivors), ~50% died perinatally | PMID: 18289368 |
|  | *Nxph1* | Fertile, premature mortality | PMID: 9856994 |
|  | *Camp (Cnlp)* | Fertile | PMID: 11719807 |
|  | *Dkk2* | Fertile | PMID: 16672341 |
|  | *Sema3e* | Fertile | PMID: 15550623 |
|  | *Apod* | Fertile | PMID: 18419796 |

### Data S1. (separate file)

DEGs_ |log₂(FoldChange)| > 1 and padj < 0.05

### Data S2. (separate file)

AMH-induced genes

## References and Notes

1. G. D. Orvis, R. R. Behringer, Cellular mechanisms of Mullerian duct formation in the mouse. Dev Biol 306, 493–504 (2007). doi:10.1016/j.ydbio.2007.03.027

2. A. Jost, Hormonal factors in the sex differentiation of the mammalian foetus. Philos Trans R Soc Lond B Biol Sci 259, 119–130 (1970).

3. A. Jost, Problems of Fetal Endocrinology - the Gonadal and Hypophyseal Hormones. Recent Prog Horm Res 8, 379–418 (1953).

4. N. Josso, WOMEN IN REPRODUCTIVE SCIENCE: Anti-Mullerian hormone: a look back and ahead. Reproduction 158, F81–F89 (2019). doi:10.1530/REP-18-0602

5. R. L. Cate, R. J. Mattaliano, C. Hession, R. Tizard, N. M. Farber, A. Cheung, E. G. Ninfa, A. Z. Frey, D. J. Gash, E. P. Chow, et al., Isolation of the bovine and human genes for Mullerian inhibiting substance and expression of the human gene in animal cells. Cell 45, 685–698 (1986). doi:10.1016/0092-8674(86)90783-x

6. O. Cohen-Haguenauer, J. Y. Picard, M. G. Mattei, S. Serero, V. C. Nguyen, M. F. de Tand, D. Guerrier, M. C. Hors-Cayla, N. Josso, J. Frezal, Mapping of the gene for anti-mullerian hormone to the short arm of human chromosome 19. Cytogenet Cell Genet 44, 2–6 (1987). doi:10.1159/000132332

7. R. D. Mullen, A. E. Ontiveros, M. M. Moses, R. R. Behringer, AMH and AMHR2 mutations: A spectrum of reproductive phenotypes across vertebrate species. Dev Biol 455, 1–9 (2019). doi:10.1016/j.ydbio.2019.07.006

8. G. D. Orvis, S. P. Jamin, K. M. Kwan, Y. Mishina, V. M. Kaartinen, S. Huang, A. B. Roberts, L. Umans, D. Huylebroeck, A. Zwijsen, D. Wang, J. F. Martin, R. R. Behringer, Functional redundancy of TGF-beta family type I receptors and receptor-Smads in mediating anti-Mullerian hormone-induced Mullerian duct regression in the mouse. Biol Reprod 78, 994–1001 (2008). doi:10.1095/biolreprod.107.066605

9. R. D. Mullen, Y. Wang, B. Liu, E. L. Moore, R. R. Behringer, Osterix functions downstream of anti-Mullerian hormone signaling to regulate Mullerian duct regression. Proc Natl Acad Sci U S A 115, 8382–8387 (2018). doi:10.1073/pnas.1721793115

10. T. Kurita, P. S. Cooke, G. R. Cunha, Epithelial-stromal tissue interaction in paramesonephric (Mullerian) epithelial differentiation. Dev Biol 240, 194–211 (2001). doi:10.1006/dbio.2001.0458

11. G. R. Cunha, Stromal induction and specification of morphogenesis and cytodifferentiation of the epithelia of the Mullerian ducts and urogenital sinus during development of the uterus and vagina in mice. J Exp Zool 196, 361–370 (1976). doi:10.1002/jez.1401960310

12. H. Du, H. S. Taylor, The Role of Hox Genes in Female Reproductive Tract Development, Adult Function, and Fertility. Cold Spring Harb Perspect Med 6, a023002 (2015). doi:10.1101/cshperspect.a023002

13. A. C. Weiss, T. Bohnenpoll, J. Kurz, P. Blank, R. Airik, T. H. Ludtke, M. J. Kleppa, L. Deuper, M. Kaiser, T. M. Mamo, R. Costa, T. von Hahn, M. O. Trowe, A. Kispert, Delayed onset of smooth muscle cell differentiation leads to hydroureter formation in mice with conditional loss of the zinc finger transcription factor gene Gata2 in the ureteric mesenchyme. J Pathol 248, 452–463 (2019). doi:10.1002/path.5270

14. M. Haugas, K. Lillevali, J. Hakanen, M. Salminen, Gata2 is required for the development of inner ear semicircular ducts and the surrounding perilymphatic space. Dev Dyn 239, 2452–2469 (2010). doi:10.1002/dvdy.22373

15. T. Moriguchi, Development and Carcinogenesis: Roles of GATA Factors in the Sympathoadrenal and Urogenital Systems. Biomedicines 9, 299 (2021). doi:10.3390/biomedicines9030299

16. Y. Zhou, K. C. Lim, K. Onodera, S. Takahashi, J. Ohta, N. Minegishi, F. Y. Tsai, S. H. Orkin, M. Yamamoto, J. D. Engel, Rescue of the embryonic lethal hematopoietic defect reveals a critical role for GATA-2 in urogenital development. EMBO J 17, 6689–6700 (1998). doi:10.1093/emboj/17.22.6689

17. M. Khandekar, N. Suzuki, J. Lewton, M. Yamamoto, J. D. Engel, Multiple, distant Gata2 enhancers specify temporally and tissue-specific patterning in the developing urogenital system. Mol Cell Biol 24, 10263–10276 (2004). doi:10.1128/MCB.24.23.10263-10276.2004

18. K. Ainoya, T. Moriguchi, S. Ohmori, T. Souma, J. Takai, M. Morita, K. J. Chandler, D. P. Mortlock, R. Shimizu, J. D. Engel, K. C. Lim, M. Yamamoto, UG4 enhancer-driven GATA-2 and bone morphogenetic protein 4 complementation remedies the CAKUT phenotype in Gata2 hypomorphic mutant mice. Mol Cell Biol 32, 2312–2322 (2012). doi:10.1128/MCB.06699-11

19. F. Y. Tsai, G. Keller, F. C. Kuo, M. Weiss, J. Chen, M. Rosenblatt, F. W. Alt, S. H. Orkin, An early haematopoietic defect in mice lacking the transcription factor GATA-2. Nature 371, 221–226 (1994). doi:10.1038/371221a0

20. Y. Lan, Q. Wang, C. E. Ovitt, R. Jiang, A unique mouse strain expressing Cre recombinase for tissue-specific analysis of gene function in palate and kidney development. Genesis 45, 618–624 (2007). doi:10.1002/dvg.20334

21. J. Chen, Y. Lan, J. A. Baek, Y. Gao, R. Jiang, Wnt/beta-catenin signaling plays an essential role in activation of odontogenic mesenchyme during early tooth development. Dev Biol 334, 174–185 (2009). doi:10.1016/j.ydbio.2009.07.015

22. Y. Lan, P. D. Kingsley, E. S. Cho, R. Jiang, Osr2, a new mouse gene related to Drosophila odd-skipped, exhibits dynamic expression patterns during craniofacial, limb, and kidney development. Mech Dev 107, 175–179 (2001). doi:10.1016/s0925-4773(01)00457-9

23. A. Ghosh, S. M. Syed, M. Kumar, T. J. Carpenter, J. M. Teixeira, N. Houairia, S. Negi, P. S. Tanwar, In Vivo Cell Fate Tracing Provides No Evidence for Mesenchymal to Epithelial Transition in Adult Fallopian Tube and Uterus. Cell Rep 31, 107631 (2020). doi:10.1016/j.celrep.2020.107631

24. J. H. Park, Y. Tanaka, N. A. Arango, L. Zhang, L. A. Benedict, M. I. Roh, P. K. Donahoe, J. M. Teixeira, Induction of WNT inhibitory factor 1 expression by Mullerian inhibiting substance/antiMullerian hormone in the Mullerian duct mesenchyme is linked to Mullerian duct regression. Dev Biol 386, 227–236 (2014). doi:10.1016/j.ydbio.2013.12.015

25. M. Kansara, M. Tsang, L. Kodjabachian, N. A. Sims, M. K. Trivett, M. Ehrich, A. Dobrovic, J. Slavin, P. F. Choong, P. J. Simmons, I. B. Dawid, D. M. Thomas, Wnt inhibitory factor 1 is epigenetically silenced in human osteosarcoma, and targeted disruption accelerates osteosarcomagenesis in mice. J Clin Invest 119, 837–851 (2009). doi:10.1172/JCI37175

26. T. Tang, L. Li, J. Tang, Y. Li, W. Y. Lin, F. Martin, D. Grant, M. Solloway, L. Parker, W. Ye, W. Forrest, N. Ghilardi, T. Oravecz, K. A. Platt, D. S. Rice, G. M. Hansen, A. Abuin, D. E. Eberhart, P. Godowski, K. H. Holt, A. Peterson, B. P. Zambrowicz, F. J. de Sauvage, A mouse knockout library for secreted and transmembrane proteins. Nat Biotechnol 28, 749–755 (2010). doi:10.1038/nbt.1644

27. D. Speidel, C. E. Bruederle, C. Enk, T. Voets, F. Varoqueaux, K. Reim, U. Becherer, F. Fornai, S. Ruggieri, Y. Holighaus, E. Weihe, D. Bruns, N. Brose, J. Rettig, CAPS1 regulates catecholamine loading of large dense-core vesicles. Neuron 46, 75–88 (2005). doi:10.1016/j.neuron.2005.02.019

28. S. Hu, R. Han, L. Chen, W. Qin, X. Xu, J. Shi, X. Zhu, M. Zhang, C. Zeng, Z. Tang, H. Bao, Z. Liu, Upregulated LRRC55 promotes BK channel activation and aggravates cell injury in podocytes. J Exp Med 218, e20192373 (2021). doi:10.1084/jem.20192373

29. N. Josso, N. Clemente, Transduction pathway of anti-Mullerian hormone, a sex-specific member of the TGF-beta family. Trends Endocrinol Metab 14, 91–97 (2003). doi:10.1016/s1043-2760(03)00005-5

30. C. S. Hill, Transcriptional Control by the SMADs. Cold Spring Harb Perspect Biol 8, a022079 (2016). doi:10.1101/cshperspect.a022079

31. Y. Zhang, T. Liu, J. Wang, B. Zou, L. Li, L. Yao, K. Chen, L. Ning, B. Wu, X. Zhao, D. Wang, Cellinker: a platform of ligand-receptor interactions for intercellular communication analysis. Bioinformatics, 2025–2032 (2021). doi:10.1093/bioinformatics/btab036

32. E. P. Gregoire, I. Stevant, A. A. Chassot, L. Martin, S. Lachambre, M. Mondin, D. G. de Rooij, S. Nef, M. C. Chaboissier, NRG1 signalling regulates the establishment of Sertoli cell stock in the mouse testis. Mol Cell Endocrinol 478, 17–31 (2018). doi:10.1016/j.mce.2018.07.004

33. D. Meyer, C. Birchmeier, Multiple essential functions of neuregulin in development. Nature 378, 386–390 (1995). doi:10.1038/378386a0

34. L. Mei, W. C. Xiong, Neuregulin 1 in neural development, synaptic plasticity and schizophrenia. Nat Rev Neurosci 9, 437–452 (2008). doi:10.1038/nrn2392

35. S. D. Harding, C. Armit, J. Armstrong, J. Brennan, Y. Cheng, B. Haggarty, D. Houghton, S. Lloyd-MacGilp, X. Pi, Y. Roochun, M. Sharghi, C. Tindal, A. P. McMahon, B. Gottesman, M. H. Little, K. Georgas, B. J. Aronow, S. S. Potter, E. W. Brunskill, E. M. Southard-Smith, C. Mendelsohn, R. A. Baldock, J. A. Davies, D. Davidson, The GUDMAP database--an online resource for genitourinary research. Development 138, 2845–2853 (2011). doi:10.1242/dev.063594

36. A. P. McMahon, B. J. Aronow, D. R. Davidson, J. A. Davies, K. W. Gaido, S. Grimmond, J. L. Lessard, M. H. Little, S. S. Potter, E. L. Wilder, P. Zhang, G. project, GUDMAP: the genitourinary developmental molecular anatomy project. J Am Soc Nephrol 19, 667–671 (2008). doi:10.1681/ASN.2007101078

37. D. N. Anbarci, J. McKey, D. S. Levic, M. Bagnat, B. Capel, Rediscovering the rete ovarii, a secreting auxiliary structure to the ovary. Elife 13, RP96662 (2025). doi:10.7554/eLife.96662

38. C. A. Rubel, S. P. Wu, L. Lin, T. Wang, R. B. Lanz, X. Li, R. Kommagani, H. L. Franco, S. A. Camper, Q. Tong, J. W. Jeong, J. P. Lydon, F. J. DeMayo, A Gata2-Dependent Transcription Network Regulates Uterine Progesterone Responsiveness and Endometrial Function. Cell Rep 17, 1414–1425 (2016). doi:10.1016/j.celrep.2016.09.093

39. T. Kurita, Normal and abnormal epithelial differentiation in the female reproductive tract. Differentiation 82, 117–126 (2011). doi:10.1016/j.diff.2011.04.008

40. G. Zhao, R. Li, Y. Cao, M. Song, P. Jiang, Q. Wu, Z. Zhou, H. Zhu, H. Wang, C. Dai, D. Liu, S. Yao, H. Lv, L. Wang, J. Dai, Y. Zhou, Y. Hu, DeltaNp63alpha-induced DUSP4/GSK3beta/SNAI1 pathway in epithelial cells drives endometrial fibrosis. Cell Death Dis 11, 449 (2020). doi:10.1038/s41419-020-2666-y

41. H. F. Stringfellow, V. J. Elliot, Endometrial metaplasia. Diagnostic Histopathology 23, 303–310 (2017). doi:10.1016/j.mpdhp.2017.05.006

42. J. W. Jeong, I. Kwak, K. Y. Lee, T. H. Kim, M. J. Large, C. L. Stewart, K. H. Kaestner, J. P. Lydon, F. J. DeMayo, Foxa2 is essential for mouse endometrial gland development and fertility. Biol Reprod 83, 396–403 (2010). doi:10.1095/biolreprod.109.083154

43. A. M. Kelleher, W. Peng, J. K. Pru, C. A. Pru, F. J. DeMayo, T. E. Spencer, Forkhead box a2 (FOXA2) is essential for uterine function and fertility. Proc Natl Acad Sci U S A 114, E1018–E1026 (2017). doi:10.1073/pnas.1618433114

44. A. M. Kelleher, F. J. DeMayo, T. E. Spencer, Uterine Glands: Developmental Biology and Functional Roles in Pregnancy. Endocr Rev 40, 1424–1445 (2019). doi:10.1210/er.2018-00281

45. M. Bouchard, A. Souabni, M. Busslinger, Tissue-specific expression of cre recombinase from the Pax8 locus. Genesis 38, 105–109 (2004). doi:10.1002/gene.20008

46. C. Mayere, V. Regard, A. Perea-Gomez, C. Bunce, Y. Neirijnck, C. Djari, N. Bellido-Carreras, P. Sararols, R. Reeves, S. Greenaway, M. Simon, P. Siggers, D. Condrea, F. Kuhne, I. Gantar, F. Tang, I. Stevant, L. Batti, N. B. Ghyselinck, D. Wilhelm, A. Greenfield, B. Capel, M. C. Chaboissier, S. Nef, Origin, specification and differentiation of a rare supporting-like lineage in the developing mouse gonad. Sci Adv 8, eabm0972 (2022). doi:10.1126/sciadv.abm0972

47. M. J. Ford, K. Harwalkar, A. S. Pacis, H. Maunsell, Y. C. Wang, D. Badescu, K. Teng, N. Yamanaka, M. Bouchard, J. Ragoussis, Y. Yamanaka, Oviduct epithelial cells constitute two developmentally distinct lineages that are spatially separated along the distal-proximal axis. Cell Rep 36, 109677 (2021). doi:10.1016/j.celrep.2021.109677

48. A. Fogarty, S. Jia, J. Wilbourne, C. DuPuis, F. Zhao, Crucial roles of mesenchymal Gata2 in murine epididymal development. Proc Natl Acad Sci U S A 122, e2507090122 (2025). doi:10.1073/pnas.2507090122

49. M. Chiga, T. Ohmori, T. Ohba, H. Katabuchi, R. Nishinakamura, Preformed Wolffian duct regulates Mullerian duct elongation independently of canonical Wnt signaling or Lhx1 expression. Int J Dev Biol 58, 663–668 (2014). doi:10.1387/ijdb.140261rn

50. A. Flesken-Nikitin, C. Q. Ralston, D. J. Fu, A. J. De Micheli, D. J. Phuong, B. A. Harlan, C. S. Ashe, A. P. Armstrong, D. W. McKellar, S. Ghuwalewala, L. H. Ellenson, J. C. Schimenti, B. D. Cosgrove, A. Y. Nikitin, Pre-ciliated tubal epithelial cells are prone to initiation of high-grade serous ovarian carcinoma. Nat Commun 15, 8641 (2024). doi:10.1038/s41467-024-52984-1

51. K. Harwalkar, N. Yamanaka, A. S. Pacis, S. Zhao, K. Teng, W. Pitman, M. Taskar, V. Lynn, A. F. Thornton, M. J. Ford, Y. Yamanaka, Aging-Associated Vacuolation of Multi-Ciliated Cells in the Distal Mouse Oviduct Reflects Unique Cell Identity and Luminal Microenvironment. Aging Cell 24, e70051 (2025). doi:10.1111/acel.70051

52. S. Jia, F. Zhao, Single-cell transcriptomic profiling of the neonatal oviduct and uterus reveals new insights into upper Mullerian duct regionalization. FASEB J 38, e23632 (2024). doi:10.1096/fj.202400303R

53. E. P. Consortium, An integrated encyclopedia of DNA elements in the human genome. Nature 489, 57–74 (2012). doi:10.1038/nature11247

54. J. Taelman, S. M. Czukiewska, I. Moustakas, Y. W. Chang, S. Hillenius, T. van der Helm, L. E. van der Meeren, H. Mei, X. Fan, S. M. Chuva de Sousa Lopes, Characterization of the human fetal gonad and reproductive tract by single-cell transcriptomics. Dev Cell 59, 529–544 e525 (2024). doi:10.1016/j.devcel.2024.01.006

55. L. Garcia-Alonso, V. Lorenzi, C. I. Mazzeo, J. P. Alves-Lopes, K. Roberts, C. Sancho-Serra, J. Engelbert, M. Mareckova, W. H. Gruhn, R. A. Botting, T. Li, B. Crespo, S. van Dongen, V. Y. Kiselev, E. Prigmore, M. Herbert, A. Moffett, A. Chedotal, O. A. Bayraktar, A. Surani, M. Haniffa, R. Vento-Tormo, Single-cell roadmap of human gonadal development. Nature 607, 540–547 (2022). doi:10.1038/s41586-022-04918-4

56. V. Lorenzi, C. Icoresi-Mazzeo, C. Cassie, N. Yayon, E. R. Ruiz-Morales, C. Sancho-Serra, R. Colligan, F. C. K. Wong, M. Mareckova, E. Tuck, K. Roberts, T. Li, M. A. Jacques, J. Ashcroft, X. He, B. Crespo, B. Cakir, S. Murray, Y. Gu, A. V. Predeus, M. Prete, I. Kelava, R. Barker, L. Garcia-Alonso, J. C. Marioni, R. Vento-Tormo, Spatiotemporal cellular map of the developing human reproductive tract. Nature 650, 428–437 (2026). doi:10.1038/s41586-025-09875-2

57. S. Allard, P. Adin, L. Gouedard, N. di Clemente, N. Josso, M. C. Orgebin-Crist, J. Y. Picard, F. Xavier, Molecular mechanisms of hormone-mediated Mullerian duct regression: involvement of beta-catenin. Development 127, 3349–3360 (2000).

58. Y. Zhan, A. Fujino, D. T. MacLaughlin, T. F. Manganaro, P. P. Szotek, N. A. Arango, J. Teixeira, P. K. Donahoe, Mullerian inhibiting substance regulates its receptor/SMAD signaling and causes mesenchymal transition of the coelomic epithelial cells early in Mullerian duct regression. Development 133, 2359–2369 (2006). doi:10.1242/dev.02383

59. A. Yamamoto, T. Omotehara, Y. Miura, T. Takada, N. Yoneda, T. Hirano, Y. Mantani, H. Kitagawa, T. Yokoyama, N. Hoshi, The mechanisms underlying the effects of AMH on Mullerian duct regression in male mice. J Vet Med Sci 80, 557–567 (2018). doi:10.1292/jvms.18-0023

60. S. Kato, T. Yokoyama, N. Okunishi, H. Narita, T. Fujikawa, Y. Kirizuki, Y. Mantani, T. Miki, N. Hoshi, Direct diffusion of anti-Mullerian hormone from both the cranial and caudal regions of the testis during early gonadal development in mice. Dev Dyn 253, 296–311 (2024). doi:10.1002/dvdy.662

61. F. M. Houghton, S. E. Adams, A. S. Rios, L. Masino, A. G. Purkiss, D. C. Briggs, F. Ledda, N. Q. McDonald, Architecture and regulation of a GDNF-GFRalpha1 synaptic adhesion assembly. Nat Commun 14, 7551 (2023). doi:10.1038/s41467-023-43148-8

62. F. Costantini, R. Shakya, GDNF/Ret signaling and the development of the kidney. Bioessays 28, 117–127 (2006). doi:10.1002/bies.20357

63. S. M. Soyal, A. Mukherjee, K. Y. Lee, J. Li, H. Li, F. J. DeMayo, J. P. Lydon, Cre-mediated recombination in cell lineages that express the progesterone receptor. Genesis 41, 58–66 (2005). doi:10.1002/gene.20098

64. J. Terakawa, V. A. Serna, D. M. Nair, S. Sato, K. Kawakami, S. Radovick, P. Maire, T. Kurita, SIX1 cooperates with RUNX1 and SMAD4 in cell fate commitment of Mullerian duct epithelium. Cell Death Differ 27, 3307–3320 (2020). doi:10.1038/s41418-020-0579-z

65. R. M. Ponnamperuma, S. M. Kirchhof, L. Trifiletti, L. M. De Luca, Ovariectomy increases squamous metaplasia of the uterine horns and survival of SENCAR mice fed a vitamin A-deficient diet. Am J Clin Nutr 70, 502–508 (1999). doi:10.1093/ajcn/70.4.502

66. Y. Yin, M. Haller, L. Goldinger, S. Bharadwaj, E. So, V. Robles-Pinos, D. Chen, L. Ma, Retinoic acid antagonizes estrogen signaling to maintain adult uterine cell fate. Proc Natl Acad Sci U S A 122, e2416089122 (2025). doi:10.1073/pnas.2416089122

67. Y. Yin, M. E. Haller, S. B. Chadchan, R. Kommagani, L. Ma, Signaling through retinoic acid receptors is essential for mammalian uterine receptivity and decidualization. JCI Insight 6, e150254 (2021). doi:10.1172/jci.insight.150254

68. Y. Zhao, S. S. Potter, Functional specificity of the Hoxa13 homeobox. Development 128, 3197–3207 (2001). doi:10.1242/dev.128.16.3197

69. A. M. Raines, M. Adam, B. Magella, S. E. Meyer, H. L. Grimes, S. K. Dey, S. S. Potter, Recombineering-based dissection of flanking and paralogous Hox gene functions in mouse reproductive tracts. Development 140, 2942–2952 (2013). doi:10.1242/dev.092569

70. K. H. Wong, H. D. Wintch, M. R. Capecchi, Hoxa11 regulates stromal cell death and proliferation during neonatal uterine development. Mol Endocrinol 18, 184–193 (2004). doi:10.1210/me.2003-0222

71. E. A. McGlade, G. G. Herrera, K. K. Stephens, S. L. W. Olsen, S. Winuthayanon, J. Guner, S. C. Hewitt, K. S. Korach, F. J. DeMayo, J. P. Lydon, D. Monsivais, W. Winuthayanon, Cell-type specific analysis of physiological action of estrogen in mouse oviducts. Faseb J 35, e21563 (2021). doi:10.1096/fj.202002747R

72. H. Q. Dinh, X. Lin, F. Abbasi, R. Nameki, M. Haro, C. E. Olingy, H. Chang, L. Hernandez, S. A. Gayther, K. N. Wright, P. J. Aspuria, B. Y. Karlan, R. I. Corona, A. Li, B. J. Rimel, M. T. Siedhoff, F. Medeiros, K. Lawrenson, Single-cell transcriptomics identifies gene expression networks driving differentiation and tumorigenesis in the human fallopian tube. Cell Rep 35, 108978 (2021). doi:10.1016/j.celrep.2021.108978

73. G. R. Cunha, J. M. Shannon, B. L. Neubauer, L. M. Sawyer, H. Fujii, O. Taguchi, L. W. Chung, Mesenchymal-epithelial interactions in sex differentiation. Hum Genet 58, 68–77 (1981). doi:10.1007/BF00284152

74. Z. Vue, G. Gonzalez, C. A. Stewart, S. Mehra, R. R. Behringer, Volumetric imaging of the developing prepubertal mouse uterine epithelium using light sheet microscopy. Mol Reprod Dev 85, 397–405 (2018). doi:10.1002/mrd.22973

75. T. E. Spencer, M. T. Lowke, K. M. Davenport, P. Dhakal, A. M. Kelleher, Single-cell insights into epithelial morphogenesis in the neonatal mouse uterus. Proc Natl Acad Sci U S A 120, e2316410120 (2023). doi:10.1073/pnas.2316410120

76. I. Passos, R. L. Britto, Diagnosis and treatment of mullerian malformations. Taiwan J Obstet Gynecol 59, 183–188 (2020). doi:10.1016/j.tjog.2020.01.003

77. A. Kobayashi, R. R. Behringer, Developmental genetics of the female reproductive tract in mammals. Nat Rev Genet 4, 969–980 (2003). doi:10.1038/nrg1225

78. A. Ludwin, I. Ludwin, M. A. Coelho Neto, C. O. Nastri, B. Bhagavath, S. R. Lindheim, W. P. Martins, Septate uterus according to ESHRE/ESGE, ASRM and CUME definitions: association with infertility and miscarriage, cost and warnings for women and healthcare systems. Ultrasound Obstet Gynecol 54, 800–814 (2019). doi:10.1002/uog.20291

79. S. Jia, J. Wilbourne, M. J. Crossen, F. Zhao, Morphogenesis of the female reproductive tract along antero-posterior and dorso-ventral axes is dependent on Amhr2+ mesenchyme in micedagger. Biol Reprod 107, 1477–1489 (2022). doi:10.1093/biolre/ioac179

80. F. Zhao, R. Li, S. Xiao, H. Diao, M. M. Viveiros, X. Song, X. Ye, Postweaning exposure to dietary zearalenone, a mycotoxin, promotes premature onset of puberty and disrupts early pregnancy events in female mice. Toxicol Sci 132, 431–442 (2013). doi:10.1093/toxsci/kfs343

81. S. Jia, F. Zhao, Ex vivo development of the entire mouse fetal reproductive tract by using microdissection and membrane-based organ culture techniques. Differentiation 123, 42–49 (2022). doi:10.1016/j.diff.2022.01.001

82. C. A. Stewart, R. R. Behringer, Mouse oviduct development. Results Probl Cell Differ 55, 247–262 (2012). doi:10.1007/978-3-642-30406-4_14

83. E. Agduhr, STUDIES ON THE STRUCTURE AND DEVELOPMENT OF THE BURSA OVARICA AND THE TUBA UTERINA IN THE MOUSE. Acta Zoologica 8, 1–133 (1927). doi:10.1111/J.1463-6395.1927.TB00649.X

